# Dynamic interactions of retroviral Gag condensates with nascent viral RNA at transcriptional burst sites: implications for genomic RNA packaging

**DOI:** 10.1101/2025.01.11.632546

**Authors:** Rebecca J. Kaddis Maldonado, Leslie J. Parent

## Abstract

Retroviruses cause significant diseases in humans and animals, including the acquired immunodeficiency syndrome and a wide range of malignancies. A crucial yet poorly understood step in the replication cycle is the recognition and selection of unspliced viral RNA (USvRNA) by the retroviral Gag protein, which binds to the psi (Ψ) packaging sequence in the 5’ leader, to package it as genomic RNA (gRNA) into nascent virions. It was previously thought that Gag initially bound gRNA in the cytoplasm. However, previous studies demonstrated that the Rous sarcoma virus (RSV) Gag protein traffics transiently through the nucleus, which is necessary for efficient gRNA packaging. These data formed a strong premise for the hypothesis that Gag selects nascent gRNA at transcription sites in the nucleus, the highest concentration of USvRNA molecules in the cell. To test this hypothesis, we used single molecule labeling and imaging techniques to visualize fluorescently-tagged, actively transcribing viral genomes, and Gag proteins in living cells. Gag foci were observed in the nucleus, transiently co-localizing with USvRNA at transcriptional burst sites, and forming viral ribonucleoprotein complexes (vRNPs). Furthermore, we found that RSV Gag interacts with Med26 and CTCF in the nucleus, suggesting a possible role for these factors in trafficking Gag to transcription sites. These results support a novel paradigm for retroviral assembly in which Gag traffics to transcriptional burst sites and interacts with nascent USvRNA through a dynamic kissing interaction to capture gRNA for incorporation into virions.

**Importance:** Retroviruses depend upon host cell transcription machinery to synthesize USvRNA, which serves as the genome selected by Gag for packaging. We previously reported that RSV Gag undergoes transient nucleocytoplasmic trafficking, which is needed for optimal genome packaging and co-localizes with USvRNA in the nucleus. Here, using live cell imaging, we found that the association of Gag with USvRNA at the transcriptional burst site is transient and dynamic. Both Gag and the RSV transcriptional burst are located near the periphery of the nucleus, which may facilitate viral RNA export. Our data also suggest that host transcription-associated factors may play a role in trafficking Gag to transcription sites.

## Introduction

RNA synthesis is coordinated with critical steps in RNA processing, including 5’ capping, splicing, polyadenylation, and 3’ cleavage, all of which occur co-transcriptionally (1-8). Many nuclear factors involved in these processes, including RNA polymerase II (RNAPII), transcription factors, and splicing machinery, coalesce into distinct nuclear foci that form dynamic biomolecular condensates (BMCs), also known transcriptional condensates (9-13). The co-transcriptional binding of these factors promotes efficient synthesis of fully-processed RNAs. The fates of cellular mRNAs are determined by specialized RNA binding proteins (RBPs) that bind during or shortly after mRNA synthesis (14-20). Spliced mRNAs are licensed for export co-transcriptionally when members of the TREX complex and Nxf1 (Tap) are recruited during splicing (15). In some cases, binding of nuclear export factors transport the mRNA to a specific subcellular location or organelle where it undergoes translation (21). Unspliced RNAs, in contrast, are typically retained in the nucleus to prevent the translation of aberrant proteins. These complex co-transcriptional processes are essential and tightly regulated, yet the mechanisms governing them are incompletely understood.

The mechanisms guiding mRNA fate during or shortly after transcription are particularly relevant for retroviruses, which cause severe immunodeficiency syndromes and cancers in humans and a wide range of animal species. Retroviruses integrate their reverse-transcribed DNA into the host cell chromosome, behaving like cellular genes transcribed by RNAPII and processed to include a 5’ cap and 3’ polyadenylated tail. Nascent retroviral RNA (vRNA) can be spliced and exported by the usual route for processed RNAs. Alternatively, the vRNA can remain unspliced and must overcome the barrier for unprocessed genes to be exported from the nucleus into the cytoplasm, where the full-length vRNA serves as (i) mRNA for synthesis of the viral structural proteins Gag and GagPol, or (ii) genomic RNA (gRNA), which is captured by Gag for packaging into new virions that propagate infection [reviewed in (22)].

The mechanism by which unspliced retroviral RNAs (USvRNAs) are sorted into mRNA or gRNA at the transcription site is incompletely understood, despite the absolute requirement for the packaging of full length vRNA to produce infectious virus particles. Recently, a novel mechanism for identifying the unspliced vRNA that serves as gRNA was proposed after finding that the retroviral Gag proteins of Rous sarcoma virus (RSV), human immunodeficiency virus type 1 (HIV-1), prototype foamy virus, murine leukemia virus, feline immunodeficiency virus, and Mason-Pfizer monkey are localized to the nucleus (23-39). In addition, both RSV and HIV-1 Gag undergo liquid-liquid phase separation to form BMCs driven by intrinsically-disordered regions (IDRs) primarily located in the NC (nucleocapsid domain) (23, 40-42). We propose that the formation of Gag BMCs permit viral condensates to remain intact while travelling through the densely-packed intracellular environment to reach the plasma membrane for budding.

To gain further mechanistic insights into the potential role of Gag nuclear trafficking in gRNA packaging, the avian retrovirus RSV was used as an experimental system because its mechanisms governing nuclear import and export are the best understood among retroviral Gag proteins. For RSV, nucleocytoplasmic trafficking of Gag is required for efficient gRNA packaging (24, 28-30). In RSV-infected cells, large, bright foci of USvRNA can be visualized in the perichromatin space using single-molecule RNA FISH (smFISH), which represent transcriptional bursts of viral RNA synthesis presumably arising at the chromosomal site of proviral integration (24). In previous studies, we found that RSV Gag localizes preferentially to the euchromatin fraction of the nucleus and co-localizes with USvRNA at transcriptional burst sites, forming viral ribonucleoprotein complexes (vRNPs) that are seen crossing the nuclear envelope during nuclear egress (24, 43).

In the present study, live-cell, time-resolved confocal imaging experiments were performed to examine the spatiotemporal interplay of Gag condensates with USvRNA at viral transcription sites to better understand the nature of this interaction. These experiments revealed the surprising finding that condensates of Gag engaged in a transient kissing interaction with nascent retroviral RNA at transcriptional burst sites, reminiscent of the interaction of RNAPII, the transcription co-factor Mediator (Med19), and actively transcribing *Sox2* mRNA, resulting in enhanced expression of the target gene (11, 13). This type of kissing interaction between a viral protein and its cognate vRNA has not been described previously, to our knowledge, therefore we sought to investigate its mechanism in more detail and examine whether the transient interaction of Gag with USvRNA at transcriptional burst sites plays a role in viral transcription regulation or gRNA packaging.

## Results

### Dynamic interaction of RSV Gag with USvRNA at transcriptional bursts

Advanced imaging approaches and single molecule labeling has revealed that large amounts of RNA are synthesized during transcription to form transcriptional bursts (37, 44). In imaging studies, transcriptional bursts appear as large, very bright nuclear RNA foci, which we previously observed in RSV-infected cells using smFISH (24). Up to now, Gag localization at viral transcription sites had only been observed in fixed cells, not allowing observation of the movement of the protein and vRNA involved in the interaction to be observed on a dynamic time scale. We were interested in examining how rapidly Gag traffics to the vRNA burst and whether the interaction is stable or transient. To gain insight into these questions, the kinetics of Gag-USvRNA interactions in the nucleus of living cells were studied in a quail fibroblast cell line (QT6), rtTA TRE RC.V8 MS2 stbl, which constitutively expresses reverse tetracycline-controlled transactivator (rtTA) and a modified RSV proviral construct controlled by a doxycycline-inducible promoter. The proviral construct incorporates 24 copies of MS2 stable stem-loops between the *nucleocapsid* (*nc)* and *protease* (*pr)* coding regions to specifically label USvRNA (45) (Figure 1A). These cells were co-transfected with pNES1-YFP-MS2-NLS, which labels USvRNA by binding to the MS2 stem-loops co-transcriptionally (46). The brightest USvRNA object(s) in each nucleus were considered to be transcriptional bursts, consistent with previous reports (11, 13). A Gag-SNAPTag fusion protein was expressed to permit single-molecule detection of Gag. In the cases of 16+ hours of doxycycline-induction, Gag-SNAPTag and NES1-YFP-MS2-NLS transfection and doxycycline treatment was performed concurrently. When cells were induced for shorter periods (∼2 hours), transfection was performed 14 hours before induction.

**Figure 1:**
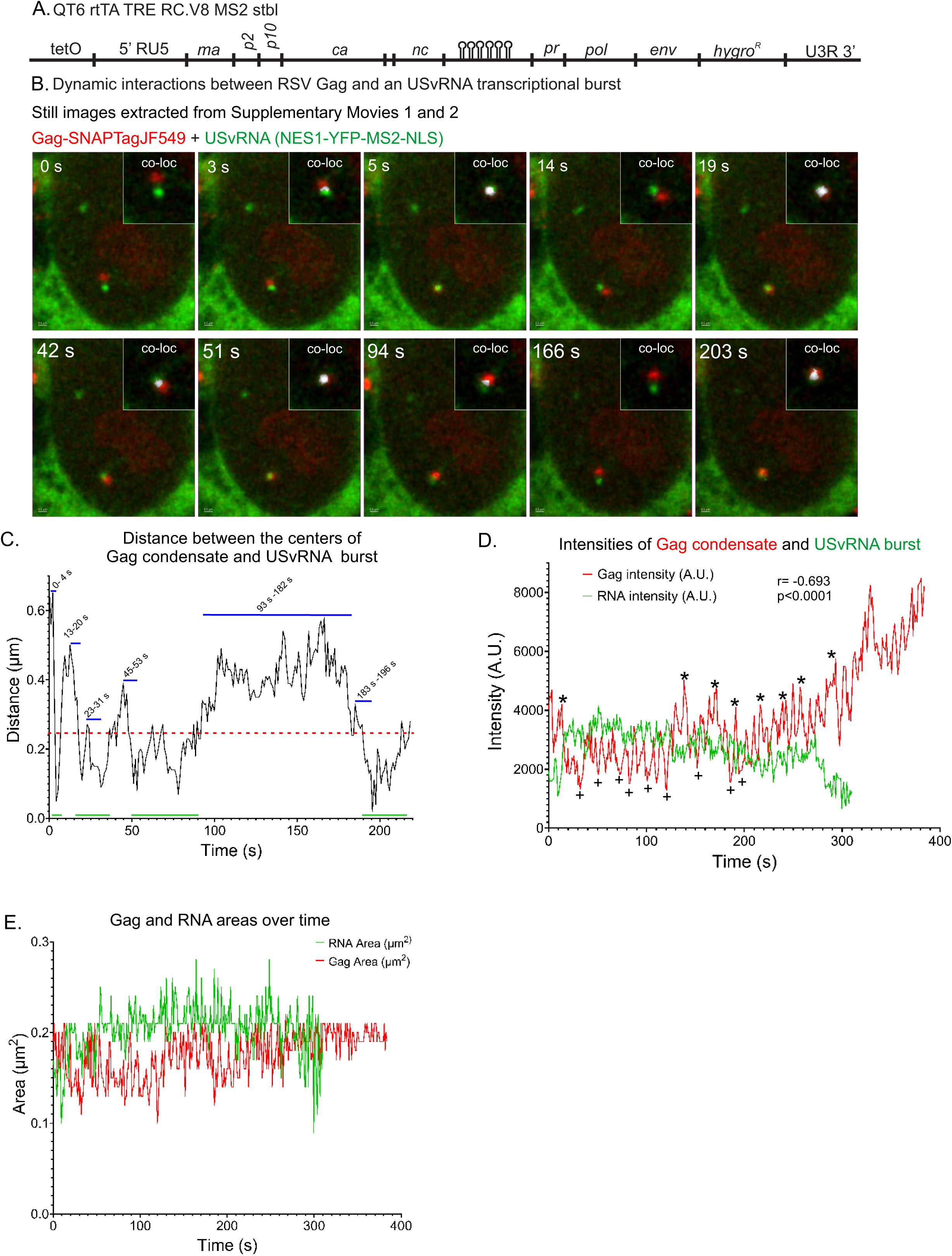
Live-cell time-lapse imaging of a Gag condensate dynamically interacting with an USvRNA transcriptional burst in a QT6 rtTA TRE RC.V8 MS2 stbl cell line. **A)** Schematic of the modified RSV provirus that was stably integrated into QT6 cells under control of a doxycycline-inducible promoter, and containing 24 copies of MS2 stable stem loops to label USvRNA. The QT6 rtTA TRE RC.V8 MS2 stbl cell line constitutively expresses rtTA. USvRNA was labeled by the MS2 coat protein fused to YFP, and containing an NLS and NES to enable MS2 to enter the nucleus while keeping nuclear background low (NES1-YFP-MS2-NLS).Cells were doxycycline-induced for ∼22 hours and imaged every 1.04 s. B) Still images from Supplementary Movie 1 show multiple instances of Gag (red) and RNA (green) undergoing transient co-localization or a “kissing” interaction. A co-localization channel was generated and overlaid with the Gag and RNA channels (white, inset, Supplementary Movie 2). C) Graph of distances between the centers of the RSV Gag condensate and the USvRNA burst. Peaks in the graph indicate the foci are apart while valleys correspond with foci within close proximity. Blue lines indicate the timepoints it takes for a peak (>0.25 µm) to dip to a valley (0.25 µm or less, green lines). Red line indicates object-based co-localization threshold (0.25 µm). D) The Gag and USvRNA are inversely correlated. r= -0.0693, p<0.0001. * indicates Gag peaks and + indicates Gag lows. E) The areas of Gag (red) and USvRNA (green) over the entire span of the movie.

After doxycycline treatment, cells were incubated with the SNAPTag ligand JF549 and imaged at ∼1 frame/s for approximately 6 minutes using confocal microscopy. Discrete condensates of Gag (red) and a large USvRNA focus (green) representing the transcriptional burst site were observed. To our surprise, these foci exhibited dynamic movement, forming kissing interactions, with Gag and vRNA foci coming together and co-localizing then moving apart multiple times over the imaging period (Figure 1B and Supplementary Movie 1). A kissing interaction was defined as co-localization of Gag and USvRNA foci at a distance of ≤0.25 µm between the centers of the Gag condensates and the vRNA bursts, based on the resolution limit of the microscope objective in the x-y plane. In some instances, however, only the edges of the USvRNA and Gag condensates overlapped, thus, the centers were further apart than the 0.25 µm distance considered as co-localized. To visualize the overlap between the edges of the Gag condensates and USvRNA, we utilized signal-based co-localization to generate a white co-localization channel to show pixels containing both fluorescent signals at the edges of condensates when the centers were ≥0.25 µm apart (Figure 1B inset, Supplementary movie 2).

To assess the temporospatial dynamics and gain insight into the underlying trafficking mechanism, particle tracking was performed to measure how rapidly co-localization and separation between the Gag condensate and vRNA burst occurred over time. Images corresponding to individual timepoints are displayed in Figure 1B, with the tracks shown in Figure 1C corresponding to the timelapse images in Supplementary Movie 1. The distances between Gag and USvRNA changed rapidly over time, with instances of separation (>0.25 µm) followed by close proximity (≤0.250 µm) in as little as 5s (Figure 1B, timespans 0-4 s and 13-20 s). The cycles of to-and-fro movement between the Gag condensate and USvRNA burst varied in duration, with the foci remaining in close proximity for ∼ 34 s (53-92 s), followed by separation (>0.250 µm) for 85 s (timepoints 93-182 s), before coming back together (183-196 s) for 30 s. In contrast, the co-localization of Med19 condensates with the *Sox2* mRNA active gene locus lasts longer, on the order of 5-10 minutes (13). These temporal differences suggest that the mechanisms that controls the Gag-USvRNA interaction differ from those regulating kissing interactions between the *Sox2* mRNA and transcriptional condensates. It is possible that the mechanisms of contact serve different purposes, for example the shorter “hit-and-run” between Gag and USvRNA could mediate gRNA packaging, whereas longer contact is needed for transcriptional condensate-mediated gene expression.

Measurements of the distances between the Gag condensate and USvRNA burst indicated that they were ≤1 µm apart at all timepoints (Supplementary Movie 1), suggesting an active mechanism maintains their close proximity. Quantitation of the fluorescence intensity of the Gag condensate demonstrated that it increased over time (Figure 1D). Simultaneously, the USvRNA fluorescence intensity decreased (Figure 1D) and eventually disappeared (300 s timepoint), possibly due to a decrease in transcriptional activity, transfer of RNA molecules from the burst to a Gag condensate, movement of the RNA outside the plane of imaging, or bleaching of the fluorophores labeling the USvRNA. Although the intensity of the Gag focus increased, the condensate area remained unchanged (ranging from 0.1-0.22 µm^2^), suggesting that the intensity increase was not caused by a change in the size of the condensate but was due to an increase in the number of Gag molecules densely packing into the condensate (Figure 1E). However, it is also possible that a change in the conformation of the Gag-SNAPTag fusion protein altered the characteristics of the fluorescence dye. Furthermore, there was an inverse correlation between the intensities of the Gag condensate and the USvRNA burst (Pearson’s correlation (r) = -0.693, p<0.0001) (Figure 1D). One possible explanation for this inverse correlation is that Gag molecules accumulate in the condensate and bind to USvRNA to form a vRNP complex, which moves away from the burst, resulting in a decrease in Gag intensity. At that point, bursting of viral transcription occurs again, with an increase in fluorescence intensity of the USvRNA focus, and the cycle repeats. The complexity of the relationship between transcriptional bursting and protein condensates has been described for cellular factors, yet remains poorly understood (9, 11, 13). Technical advances in super-resolution imaging or other biophysical techniques will be needed to dissect how and why newly transcribed USvRNA and Gag engage in such complex choreography.

Quantitative analysis of a second live-cell experiment demonstrated numerous to-and-fro movements between a Gag condensate and an USvRNA transcription burst site (Figure 2A-C; Supplementary Movie 3). Particle tracking of the Gag condensate and USvRNA burst site indicated that they remained within close proximity (0.7 µm) of one another during the 3-minute duration of imaging (Figure 2B). The Gag condensate moved towards the transcriptional burst and underwent co-localization in ∼51 s. The kissing interaction was initially brief, and the distance between the Gag condensate and USvRNA then fluctuated from near to far between timepoints 56-102 s. Following that initial contact, there was a long period of co-localization lasting 28 s (timepoints 102-129 s) followed by a long separation (43 s; timepoints 129-172 s) and then a brief period of co-localization. Similar to the data shown in Figure 1D, the fluorescence intensity of the Gag condensate signal was inversely correlated with the USvRNA intensity (r = -0.454, p<0.0001, analyzed from 1-200s, Figure 2C).

**Figure 2:**
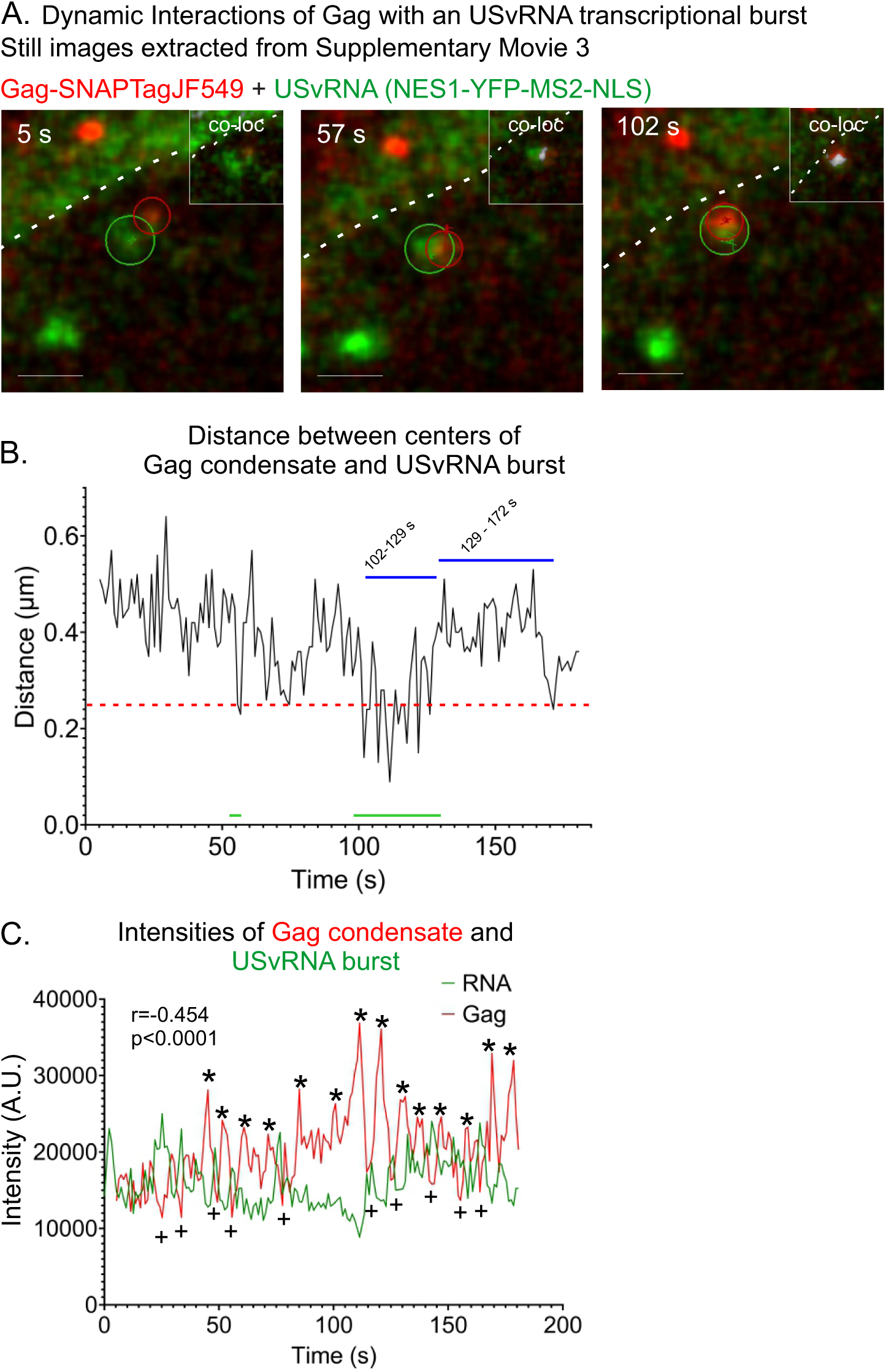
Dynamic interactions of Gag with an USvRNA transcriptional burst observed via of live-cell time-lapse imaging of QT6 rtTA TRE RC.V8 MS2 stbl cell line. **A)** Stills correlating to Supplementary Movie 3. This cell was induced for ∼16 hours and imaged every second. A co-localization channel was generated from the Gag and RNA channels and overlaid (white, inset). Scale bar = 1 µm. **B)** Graph of distances between the centers of the RSV Gag condensate and the USvRNA burst. The distance between Gag and the burst remains within 1 µm. Blue lines indicate the timepoints it takes for a peak (>0.25 µm) to dip to a valley (0.25 µm or less, green lines). Red line indicates object-based co-localization threshold (0.25 µm). **C)** The Gag and USvRNA intensities are inversely correlated. R= -0.454, p<0.0001. Gag peaks are marked by * and lows are marked by +.

At a different time point in the same cell, we observed multiple Gag condensates near two separate transcriptional bursts (Figure 3A, Supplementary Movie 4). This set of images indicated that more than one Gag condensate can enter the nucleus and make transient contact with more than one USvRNA burst sites. Two of the Gag condensates were tracked and even though there were two USvRNA bursts, the Gag condensates appeared to favor the burst on the left over the burst on the right. We have observed this phenomenon previously in acutely-infected fixed cells where Gag was co-localized with one burst but not the other (24). It is feasible that the bursts are at different stages of transcription and Gag preferentially co-localizes with one stage over the other. Another possibility is that the nuclear topology blocks access of the Gag condensate to one of the vRNA transcription sites due to its location on a particular euchromatin loop or the local environment of the proviral integration site. Further studies will be needed to investigate these possibilities.

**Figure 3:**
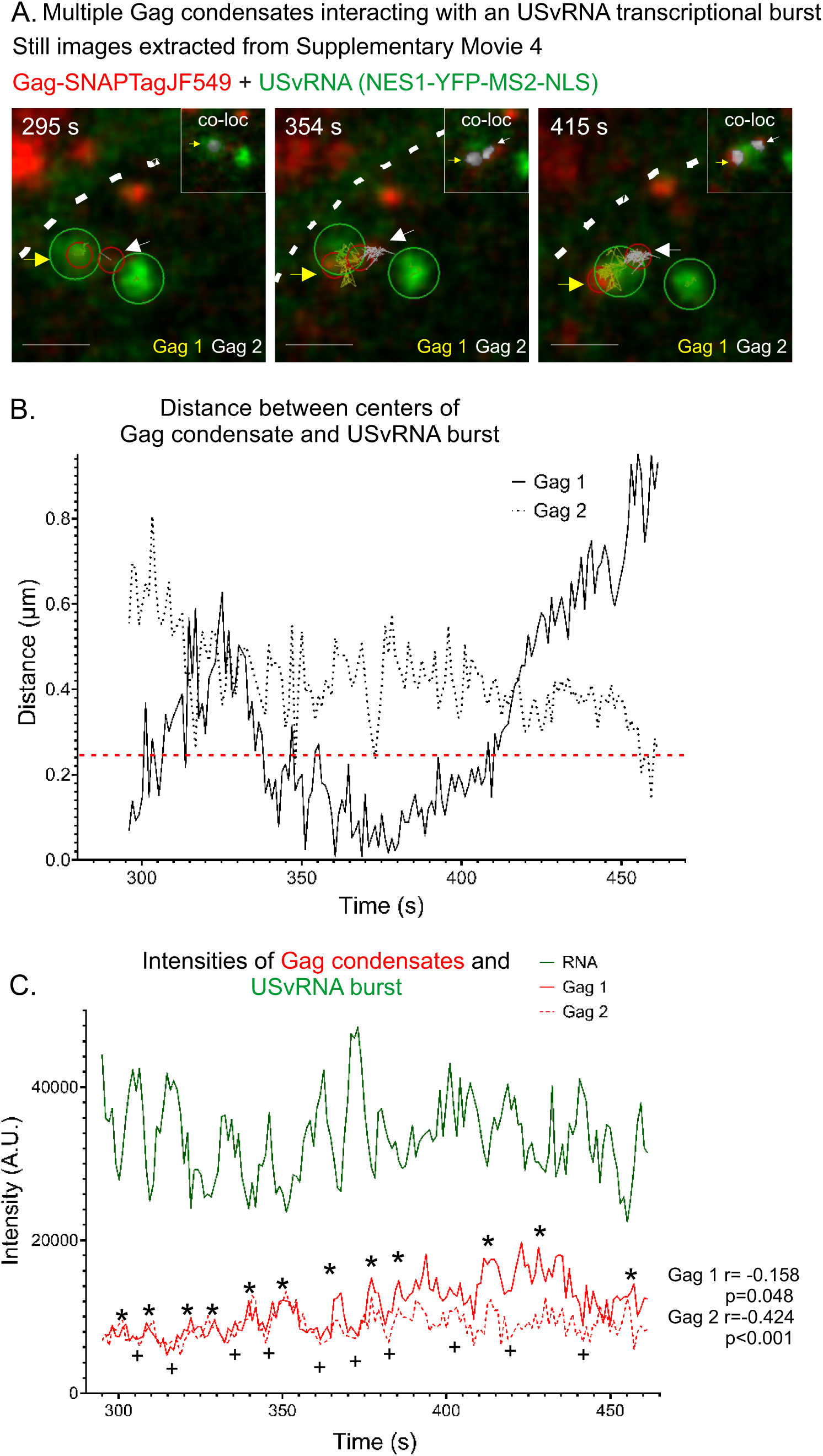
Multiple Gag condensates interacting with an USvRNA transcriptional burst observed via of live-cell time-lapse imaging of QT6 rtTA TRE RC.V8 MS2 stbl cell line. **A)** Stills from Supplementary Movie 4 (16 hours post-induction, ∼1 frame/second) showing multiple Gag condensates at two bursts. Gag 1 is marked with a yellow arrow and track. Gag 2 is marked by a white arrow and track. A co-localization channel was generated from the Gag and RNA channels and overlaid (white, inset). Scale bar = 1 µm. **B)** Graph of distances between the centers of the RSV Gag condensates and the USvRNA burst. Both Gag condensates remained within 1 µm of the burst. Red line indicates object-based co-localization threshold (0.25 µm). **C)** The intensities of both Gag condensates are once again inversely correlated with that of the USvRNA burst. Gag 1: r= -0.158, p=0.048. Gag 2: r= -0.424, p<0.001. Gag peaks are marked by * and lows are marked by +. The nuclear rim is marked by the white dotted line.

Live cell particle tracking (Figure 3A and B; Supplementary Movie 4) revealed that condensate #1 (Gag 1,yellow arrow and tracking) was co-localized with the USvRNA burst for ∼75 s, and as it moved away from the vRNA, Gag condensate #2 (Gag 2, white arrow and tracking) moved toward the USvRNA burst and became co-localized. Consistent with Figures 1D and 2C, the intensities for Gag condensates #1 and #2 were inversely correlated to the intensity of the USvRNA transcriptional burst throughout the course of the real time imaging period shown in Figure 3C (Gag 1 intensity to USvRNA intensity: r = -0.158, p=0.048; Gag 2 intensity to USvRNA intensity: r = -0.424, p<0.001).

To determine whether Gag-USvRNA kissing interactions could be observed at shorter periods after doxycycline induction, cells were induced for only two hours before imaging (Figure 4A, Supplementary Movie 5). A Gag focus initially visualized in the cytoplasm (white arrowhead, Figure 4A; 0 s) subsequently crossed into the nucleus (dashed white line), moving toward the burst of USvRNA transcription. The elapsed time from when the Gag condensate entered the nucleus and trafficked to the transcription site was ∼173 s. Once the Gag condensate entered the nucleus, it took ∼137 s to co-localize (≤0.25 µm) with the USvRNA burst and displayed a “hit-and-run” interaction with the burst over a period of 30 s (310 s-339 s, white channel, Figure 4A inset). The USvRNA burst was positioned near the nuclear rim, as reported for actively transcribing genes (47), near the point where Gag entered the nucleus, which could explain how the Gag condensate trafficked to the transcriptional burst with rapid kinetics.

**Figure 4:**
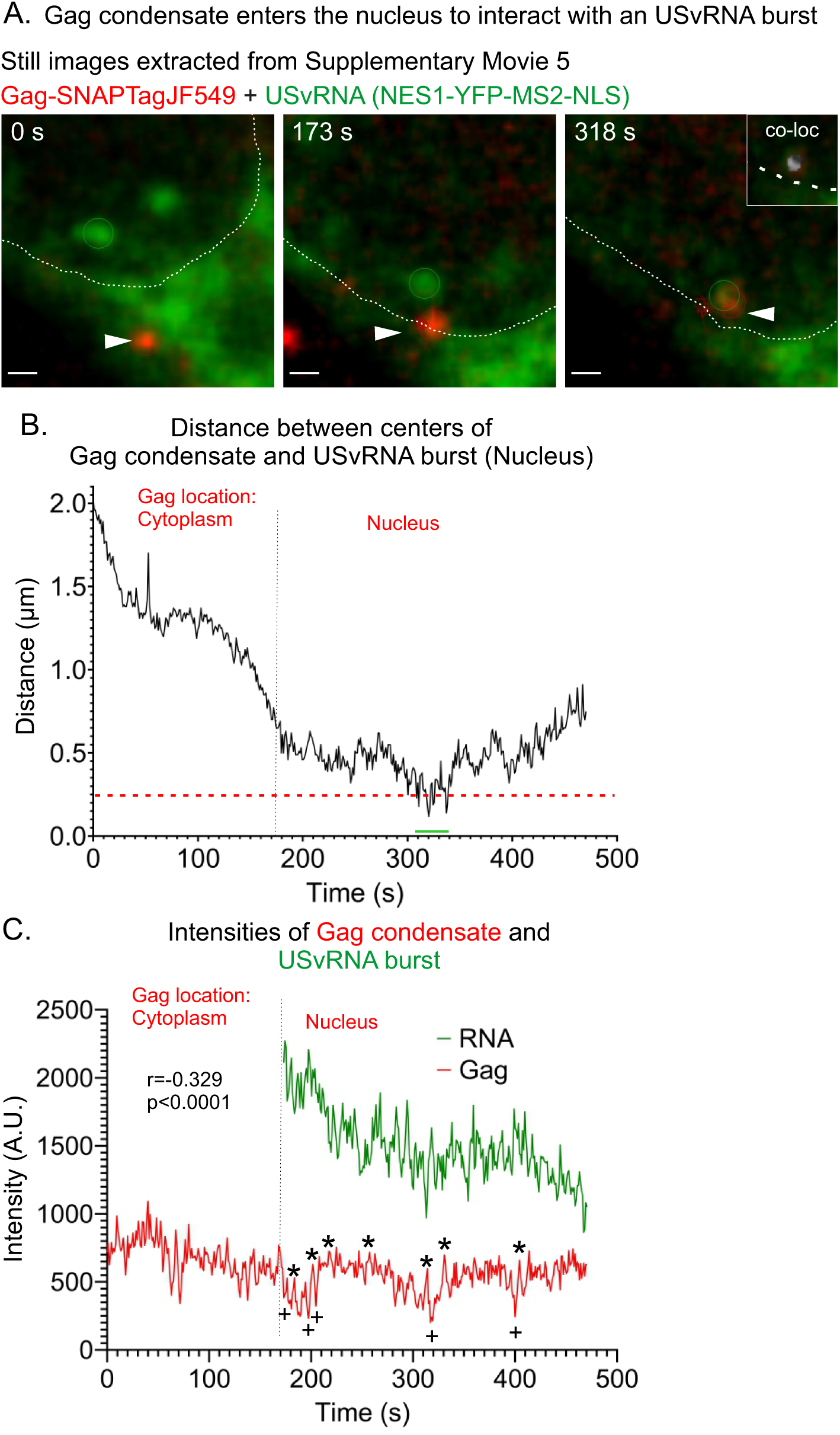
Gag condensate enters the nucleus to interact with an USvRNA burst observed via live-cell time-lapse imaging of QT6 rtTA TRE RC.V8 MS2 stbl cell line. **A)** Stills correlating to Supplementary Movie 5 of a cell 2 hours post induction (imaged ∼ 1 frame/ second) in which Gag (red) traffics from the cytoplasm into the nucleus (white outline) to the USvRNA burst (green) before undergoing co-localization. Scale bar = 0.5 µm. A co-localization channel was generated and overlaid with the gag and RNA channels (white, inset, t=318 s). **B)** Graph of distances between the centers of the RSV Gag condensate and the USvRNA burst measured over time. The red text indicates the location of Gag during those time points and the black dotted line indicates when Gag crosses into a new compartment. Red line indicates object-based co-localization threshold (0.25 µm). **C)** Intensities of Gag and USvRNA condensates over time. USvRNA intensity is only being shown once Gag enters the nucleus. The Gag and USvRNA intensities are inversely correlated. r= -0.329, p<0.0001. Gag peaks are marked by * and lows are marked by +.

From the time the Gag condensate entered the nucleus, it remained in close proximity with the USvRNA burst (≤0.9 µm; Figure 4B) for over 5 minutes, until the end of the imaging time. The intensity of the Gag signal remained constant from its position in the cytoplasm throughout its residence in the nucleus (Figure 4C). However, the RNA signal diminished over time, possibly due to a decrease in transcriptional activity, movement out of the imaging plane, or bleaching of the fluorescence signal from imaging (Figure 4C). Additionally, the decrease in RNA signal over time could be due to Gag molecules forming complexes with USvRNA molecules and carrying it away from the burst, that we are unable to see due to the limited resolution of confocal microscopy. The intensities of the Gag and USvRNA signals were inversely correlated, as seen in each of the previous episodes (r = -0.329, p<0.0001). Due to the dynamic nature of Gag nuclear trafficking, this event was difficult to capture. Although we have observed many Gag foci that enter the nucleus, this is the first time we have seen Gag enter the nucleus to co-localize with the USvRNA burst.

We previously reported that Gag interacts with the nuclear export protein CRM1 to mediate its nuclear egress (29, 48) and observed Gag-vRNA complexes located near the nuclear envelope cross into the cytoplasm using a transient transfection system (24). To determine whether a similar export of the vRNP complex could be imaged using the stable cell line shown in Figure 5A (Figure 1), we obtained live cell images showing the Gag-USvRNA complex undergoing nucleocytoplasmic trafficking (Supplementary Movie 6). Still images extracted from this live cell movie (Figure 5A) show a Gag condensate co-localized with a focus of USvRNA, with the vRNP complex crossing the nuclear rim (372 s) into the cytoplasm. This USvRNA focus was not defined as a transcriptional burst site because it was not the brightest USvRNA focus in the nucleus, and it was mobile rather than stationary. A co-localization channel (white) illustrates the overlapping Gag-USvRNA signals in the vRNP complex (Figure 5A inset, upper right corner, white signal and track; see also Supplementary Movie 7) as it was undergoing nuclear egress. The co-localized vRNP moved in a to-and-fro fashion along the nuclear edge several times during the movie, and after arriving in the cytoplasm, the Gag-USvRNA foci remained stably associated (distance ≤0.25 µm) (Figure 5B).

**Figure 5:**
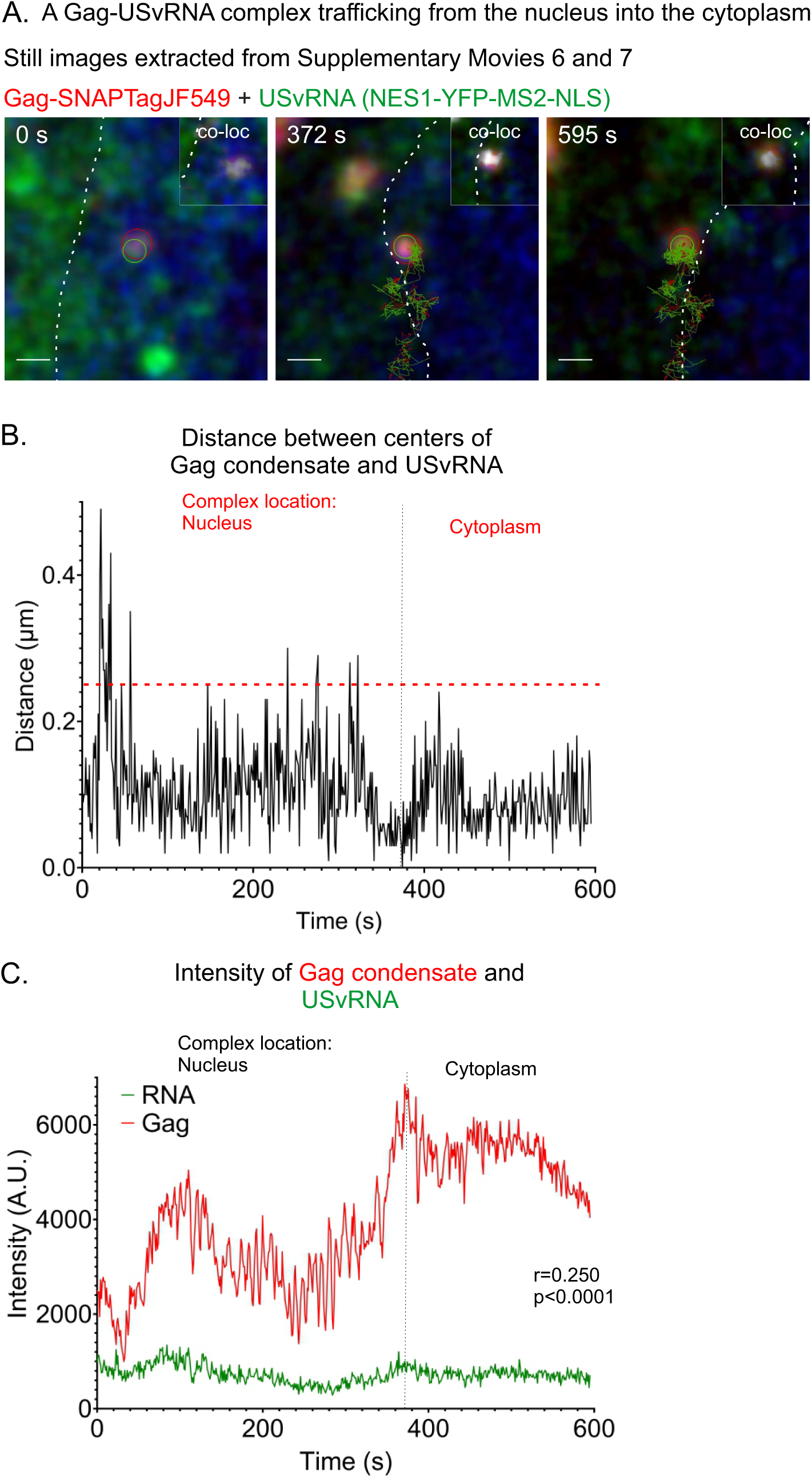
Instance of a Gag-USvRNP complex trafficking out of the nucleus observed via live-cell time-lapse imaging of QT6 rtTA TRE RC.V8 MS2 stbl cell line. **A)** Stills correlating the Supplementary Movies 6 (overlay) and 7 (co-localization channel) showing a vRNP composed of Gag and USvRNA trafficking through the nucleus into the cytoplasm. A co-localization channel was generated and overlaid with the gag and RNA channels (white, inset). **B)** Graph of distances between the centers of the RSV Gag condensate and the USvRNA focus measured over time. The red text indicates the location of the Gag-USvRNA complex during those time points and the black dotted line indicates when the complex crosses into a new compartment. Red line indicates object-based co-localization threshold (0.25 µm).The condensates remain within close proximity (<0.5 µm). **C)** The Gag and USvRNA intensities are positively correlated. r= 0.25, p<0.0001. The nuclear rim is marked by the white dotted line and/ or NucSpot650 (blue). Gag peaks are marked by * and lows are marked by +.

When we measured the intensity of the Gag condensate in Figure 5A, we found that it changed in an undulating pattern over time (Figure 5C), suggesting that additional Gag molecules were joining and leaving the condensate. Alternatively, the Gag condensate could have moved in and out of the imaging plane or the conformation of the Gag fusion protein could have changed, resulting in alteration of the quantum characteristics of the SNAP Tag ligand. In contrast to the previous Gag condensate in Figure 1, the Gag and USvRNA signals remained co-localized and the intensities were positively correlated (Figure 5C; r=0.250, p<0.0001). The observation that this Gag-USvRNA complex moved out of the nucleus and into the cytoplasm, suggests that this vRNP represents an early step in the journey of the packaging condensate moving toward the cell membrane for release as a virus particle. Due to the dynamic nature of RSV Gag nuclear trafficking and the low abundance of Gag present in the nucleus under steady-state conditions, it is challenging to capture Gag entering or exiting the nucleus and localizing transiently at transcription sites. Very few kissing events were captured under steady state conditions (7 cells with kissing interactions were captured from 9 replicates for 16-22 hour dox induction) and 4 cells from 6 replicates (for 2 hour dox induction).

To observe this transient interaction more readily, cells were transfected as above with the addition of dominant negative mutant nucleoporin proteins consisting of the FG repeats derived from Nup98 (NP98) or Nup214 (NP214)(49) (Figure 6). From our previous work (34), we knew that NP98 and NP214 trapped Gag in the nucleus due to the role of each of the Nups in CRM1-mediated nuclear export of Gag. This method of blocking Gag nuclear export was preferable to (i) mutating the Gag nuclear export signal in p10, since it allowed us study the wild-type Gag protein, or (ii) treating with the CRM1 inhibitor leptomycin B (LMB), which blocks the nuclear export of many cellular proteins as well as Gag (34). Cells were transfected with Gag-SNAPTag, NES1-YFP-MS2-NLS, NP98 or NP214 plasmids, and viral gene expression was dox-induced for two hours before imaging. Cells were imaged every ∼1 frame/s for 5 minutes through a single plane in the center of the nucleus using focus control to minimize drifting. Due to the dominant negative activity of the NP mutants, the Gag protein, USvRNA, and unbound NES1-YFP-MS2-NLS coat protein all accumulated in the nucleus, making it possible to capture several instances of kissing between Gag and USvRNA located at transcriptional bursts (Supplementary Movie 8). Compared to live-cell experiments in the absence of NP mutants, we observed kissing events in 10 cells from 2 replicates with NP98 and 9 cells from 2 replicates with NP214. In still images extracted from Supplementary Movie 8 (Figure 6A) in cells expressing NP98, the kissing interaction between Gag (red) and USvRNA burst (green) was clearly observed, and there was co-localization (white) (Figure 6A) of the condensates (Supplementary Movie 8). Interestingly, the centers of each condensate remained >0.25 µm apart (Figure 6B) for most of the imaging time, and Gag remained within 1 µm of the burst.

**Figure 6:**
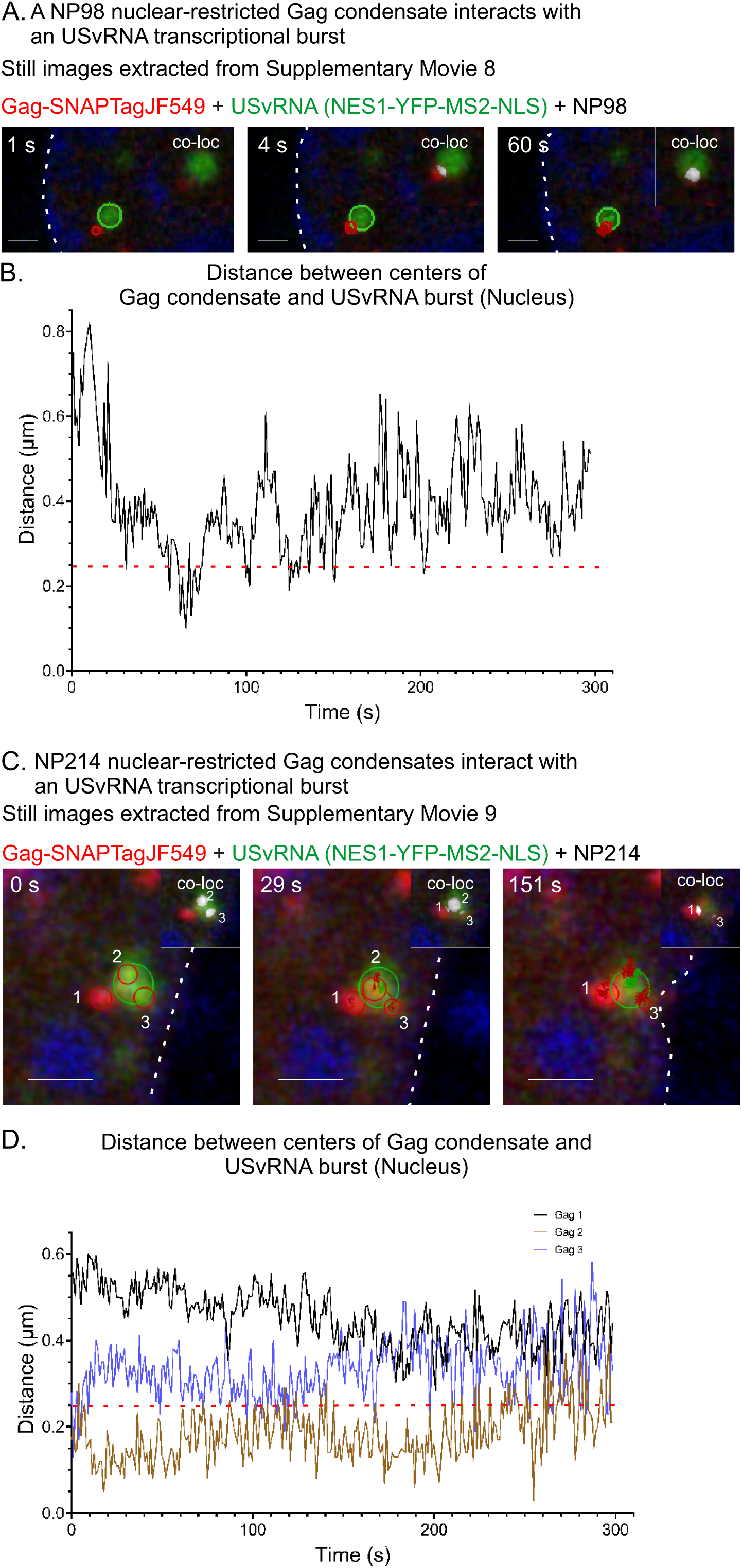
Instances of co-localization between Gag and USvRNA observed via of live-cell time-lapse imaging of QT6 rtTA TRE RC.V8 MS2 stbl cell line in the presence of dominant-negative NP proteins. QT6 rtTA TRE RC.V8 MS2 stbl cells induced for ∼2 hours (imaged ∼ 1 frame/ second) in which Gag (red) and USvRNA labeled via NES1-YFP-MS2-NLS (green) are trapped in the nucleus (white outline, Blue-DRAQ5) via dominant negative FG repeats from either Nup98 (A-C, NP98) or Nup214 (D-F, NP214). **A)** Stills correlating to Supplementary Movie 8 of a Gag condensate interacting with the USvRNA burst (green) in which Gag and USvRNA is unable to leave the nucleus in the presence of NP98. A co-localization channel was generated and overlaid with the gag and RNA channels (white, inset). Scale bar = 1 µm. **B)** Graph of distances between the centers of the RSV Gag condensate and the USvRNA burst. The distance between Gag and the burst remains within 1 µm. Red line indicates object-based co-localization threshold (0.25 µm). **C)** Stills correlating to Supplementary Movie 9 of three Gag condensates interacting with the USvRNA burst (green) in which Gag and USvRNA is unable to leave the nucleus in the presence of NP214. A co-localization channel was generated and overlaid with the Gag and RNA channels (white, inset). Scale bar = 1 µm. The Gag condensates present in each panel are numbered. **D)** Graph of distances between the centers of the RSV Gag condensates and the USvRNA burst. All Gag condensates remained within 1 µm of the burst. Red line indicates object-based co-localization threshold (0.25 µm).

Similarly, in the presence of NP214 (Figure 6C, Supplementary Movie 9), three separate Gag condensates (marked as 1, 2, and 3) were observed linked to the same burst, and object-based co-localization was used to measure the distance between that the centers of each Gag condensate and the vRNA burst. Gag 1 (black line, Figure 6D) remained >0.25 µm separated from the burst; Gag 2 remained co-localized with the burst (<0.25 µm); and Gag 3 was not colocalized. However, using the co-localization channel (white, Figure 6C inset, Supplementary Movie 9), there was clearly overlap between the edges of each of the Gag 1, 2, and 3 condensates with the vRNA burst. In cells presented in Figure 6, the Gag condensates appear more restricted in their movements compared to those without NP expression (Figures 1-3), suggesting that the NP98 and NP214 mutants constrained the movement of Gag, as seen in our previous study of a Gag-NES mutant that exhibited obstructed diffusion (37). An explanation for this finding may be found in a recent report showing that yeast CRM1 brings chromatin-bound transcription factors toward the nuclear pore to facilitate RNA transcription at the nuclear periphery (50). Since CRM1 interacts with the FG repeats in Nup98 and Nup214 to facilitate nuclear export, it is possible that NP98 and NP214 (50-52) prevent that interaction, resulting in tethering Gag at transcription sites.

Due to the non-specific nucleic acid binding ability of the Gag NC domain, we asked whether the co-localization between Gag and newly transcribed RNA also occurred with non-viral RNAs. For this experiment we expressed a Gag nuclear export mutant (Gag.L219A-CFP) which is trapped in the nucleus and forms discrete puncta (48). Previously, we found that 82±3% of USvRNA foci co-localized with Gag.L219A foci (24). To look at non-viral RNA interacting with Gag.L219A, cells were either co-transfected with pSL-MS2-24x, which expresses a non-viral RNA encoding 24 copies of MS2 stem loops, and pMS2-YFP-NLS coat protein (45, 46, 53), or pulsed with 5-fluorouridine (5FU) for 10 minutes to label nascent RNAs and labeled via immunofluorescence (Supplementary Figure 1A and B). Cells were fixed, single confocal planes were imaged, and co-localization analysis was performed via MatLab (24). In cells expressing pSL-MS2-24x, 18±4% of RNA foci co-localized with Gag.L219A (***p<0.0001). When cells were pulsed with 5FU, 10±4 (***p<0.0001) of 5FU foci co-localized with Gag.L219A foci. Compared to USvRNA with Gag.L219A (24), co-localization of non-viral RNA was significant lower, suggesting specific binding of Gag to USvRNA (Supplementary Figure 1C). However, the low-level co-localization of Gag with non-viral RNA suggests that Gag may traffic to active transcription sites and “sample” nascent RNAs in search of USvRNA at the burst.

### RSV Gag condensates co-localized with nascent USvRNA at viral transcription sites

To rigorously test whether RSV Gag was binding to nascent USvRNA at the viral transcription site, QT6 cells expressing RC.V8 Gag-SNAPTag and MS2 stem-loops (Figure 7A) were dox-induced for 24 hours and incubated with 5-Ethynyl Uridine (EU). Nascent RNA was pulse-labeled with EU for 10 minutes to label viral and cellular RNAs. Click chemistry detected cellular RNA, smFISH was used to specifically detect USvRNA, and Gag was detected via immunofluorescence. Cells were imaged with confocal microscopy and three-dimensional cross-sections were generated from Z-stacks (Figure 7B). Three-way signal-based co-localization (yellow) analysis revealed that the USvRNA (green), Gag (red), and EU (gray) were co-localized in the nucleus (dashed white line). Figure 7C shows an enlargement of the area of interest to illustrate the 3-way co-localization, indicating that nuclear Gag was associated with newly transcribed USvRNA at active transcription sites.

**Figure 7:**
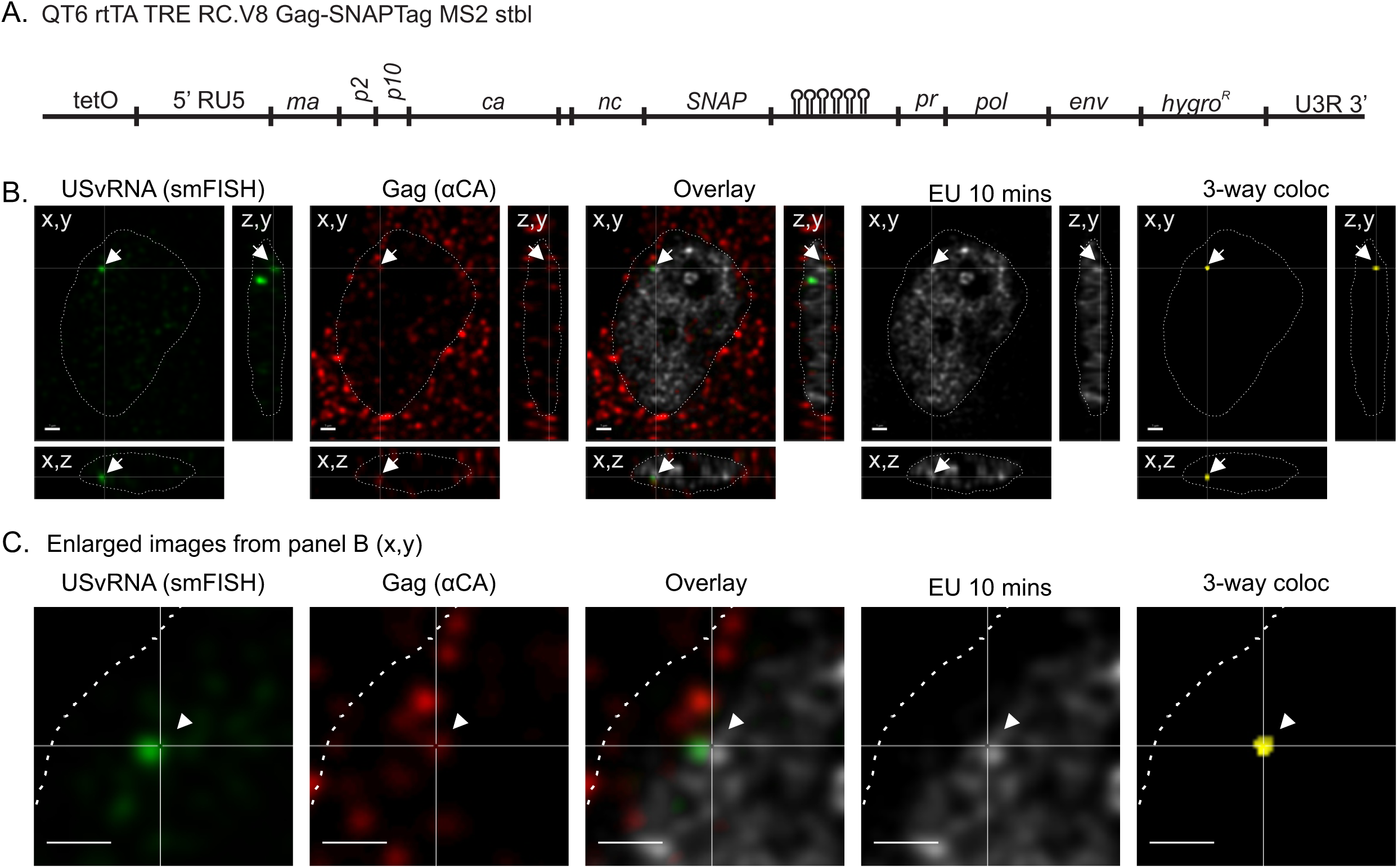
Gag co-localizes with nascent USvRNA at transcriptional bursts. **A)** QT6 rtTA TRE RC.V8 Gag-SNAPTag MS2 stbl cells constitutively express rtTA, and contain a stably integrated, modified RSV provirus that is under control of a doxycycline-inducible promoter, expresses a Gag-SNAPTag fusion protein, and contains 24 copies of MS2 stable stem loops to label USvRNA. **B)** QT6 rtTA TRE RC.V8 Gag-SNAPTag MS2 stbl cells were dox-induced for 24 hours. In the last 10 minutes, cells were pulse labeled with EU. USvRNA was labeled via smFISH, Gag labeled via immunofluorescence, and EU labeled-RNAs were subjected to Click-chemistry to label them with Alexa 488. Z-stacks of cells were imaged via confocal microscopy and used to generate cross-sections. A burst of USvRNA (green), co-localized (white arrow) with Gag (red), and EU labeling (grey) in the nucleus (DAPI-blue, white outline). Three-way co-localization (yellow) was conducted to confirm this finding. Scale bar = 1 µm. **C)** An enlargement of the image presented in B. Scale bar = 1 µm. N=3 replicates.

### RSV USvRNA transcriptional bursts were located within 1 µm of nuclear edge

Given that the HIV-1 provirus preferentially integrates within 1 µm of the nuclear envelope (35, 54), we performed confocal imaging experiments to determine the location of RSV transcriptional bursts in infected cells. Although the proviral DNA was not directly labeled, we expect the USvRNA transcriptional burst site is at the same location as the integrated provirus because RNA polymerase II uses the integrated provirus as the template for transcription. All of the USvRNA bursts (n=95, Supplementary Table 1) were <1.0 µm (mean distance = 0.31 µm ± 0.03 µm) from the edge of the nucleus (defined by DAPI) in three dimensions, indicating that like HIV-1, RSV integrates close to the nuclear rim (Figures 8A, 9A, and Supplementary Table 1). Similarly, nearly all of the Gag condensates (91.8%, n=1313 out of 1331) in the nucleus were located within 1 µm of the nuclear periphery (mean distance = 0.14 µm ± 6.81x10^-3^ µm; Figures 8B and 9A, Supplementary Table 2). Together, these data indicate that both Gag and the USvRNA are positioned near the edge of the nucleus, so Gag condensates do not need to travel far into the interior of the nucleoplasm in search of the USvRNA burst. Future experiments will examine whether nuclear Gag condensates located farther inside the nucleus could be performing other functions, such as altering chromatin organization, splicing, or influencing other cellular processes (43).

**Figure 8:**
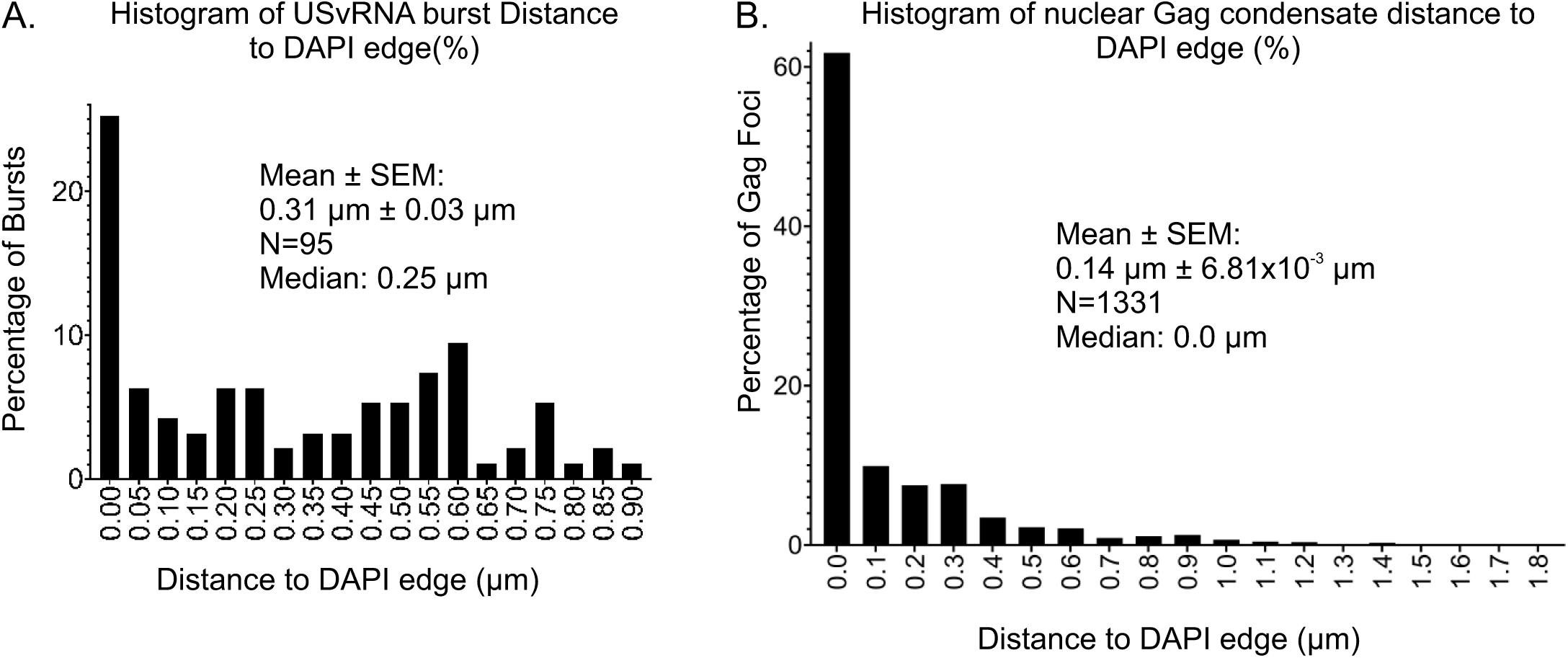
USvRNA bursts and nuclear Gag localize near the nuclear rim. **A)** All bursts were within 1 µm of the nuclear rim (as marked by DAPI in three-dimensions), with an average of 0.31 µm ± 0.03 µm. **B)** 91.8% of Gag foci are present within 1 µm of the nuclear boundary, at an average distance of 0.14 µm ± 6.81x10^-3^ µm. n=42 cells from four replicates.

**Figure 9:**
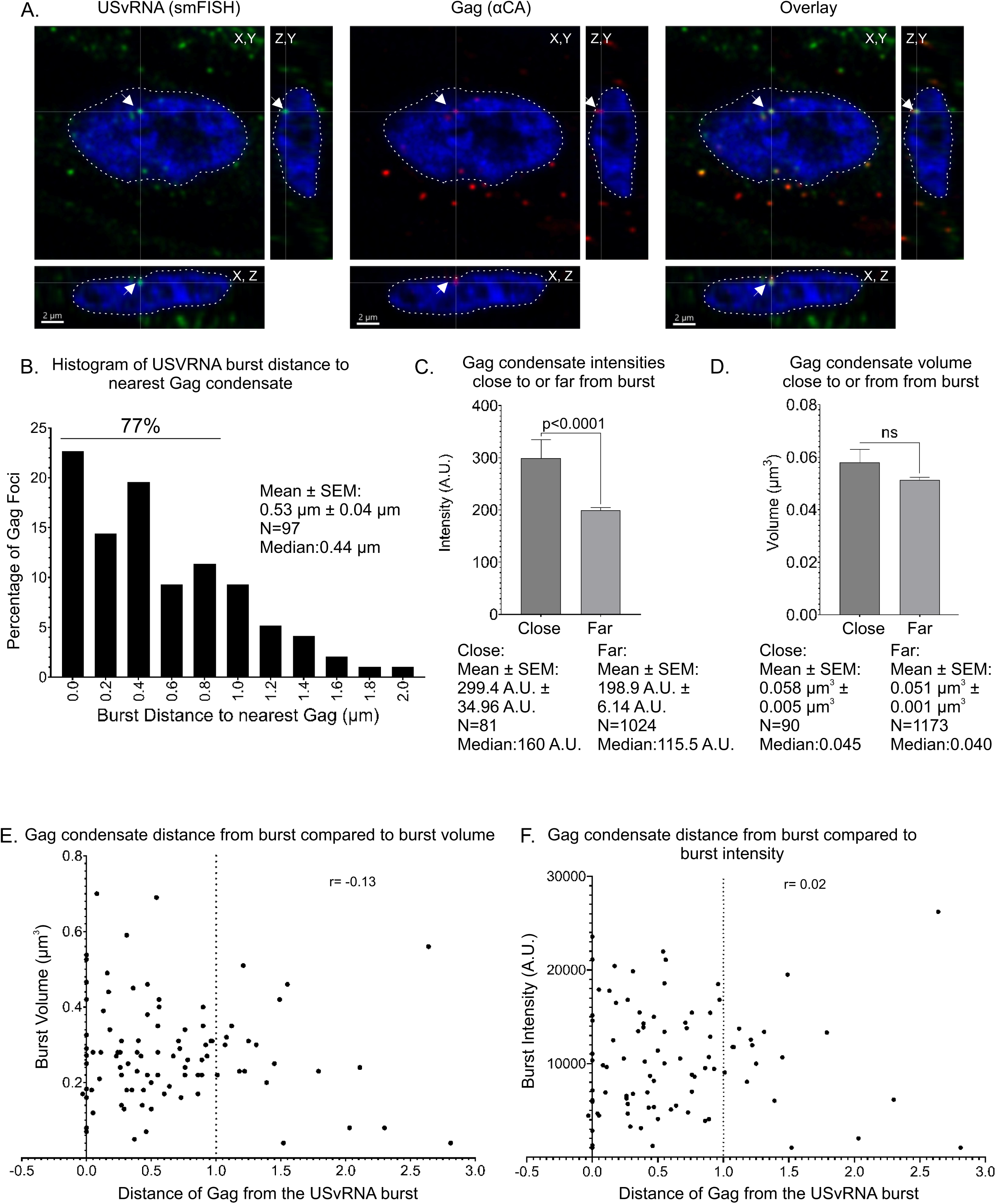
Gag localizes in close proximity of transcriptional bursts in chronically infected cells. **A)** QT6 cells chronically infected with RSV with subjected to simultaneous smFISH to label USvRNA (green) and immunofluorescence to label Gag (red). Cells were imaged via confocal microscopy, and Z-stacks were used to generate cross-sections. A burst of USvRNA (green) co-localizes with Gag (red) in the nucleus (DAPI-blue, white outline). Scale bar = 2 µm. **B)** Histogram of USvRNA burst distance to nearest Gag focus. 77% of Gag nuclear foci are localized within 1 µm of the USvRNA burst with an average distance of 0.536 µm. (N=97 bursts). **C)** The average intensity of Gag nearest the burst (299.4 A.U. ± 34.96) was statistically significantly higher (****p<0.0001) than that of Gag foci away from the burst (198.9 A.U. ± 6.140 A.U.) while there was no significant difference in the volumes between Gag closest compared to those away from the burst **D)**. **E)** There is very low correlation between Gag distance from the burst and burst volume (r = -0.13) nor **F)** burst intensity (r = 0.02). Vertical dotted line indicates 1 µm distance from the burst. n=42 cells from four replicates.

### Complex morphology of USvRNA transcriptional bursts revealed by STED microscopy

The high intensity of transcriptional bursts is attributed to the large quantity of nascent RNA being produced, with individual RNA molecules undergoing different stages of transcription along with co-transcriptional RNA processing steps (11, 13, 55, 56). To elucidate more structural detail of the RSV USvRNA bursts in infected cells, we used super-resolution STED microscopy (green) and compared that method to images obtained by confocal microscopy (red). The smFISH probes were complementary to the RSV intronic sequence to specifically detect USvRNA, and the inner leaflet of the nuclear membrane was outlined with Sun1-Venus (blue)(Supplementary Figure 2).

In a single z-slice (Supplementary Figure 2A), the signals from the confocal and STED images overlapped, as expected, but there was more detail seen in the STED images (Supplementary Figure 2A). A three-dimensional reconstruction was performed with orthogonal clipping planes of surface renderings of the transcriptional burst showing that the contour of the RNA signal looked smooth and indistinct in the confocal images, whereas using STED, the burst surfaces appeared sharper, with multiple connected nodes visualized (Supplementary Figure 2B). These nodes could be regions of high transcriptional activity during bursting, as additional molecules of RNAPII are recruited to the integrated proviral DNA (56). To show more detail, magnified images of two different bursts are shown in panels C and D. In both cases, multiple foci of RNA appeared to be connected, forming a complex structure, which may represent RNA emanating from clustered transcriptional condensates (57, 58). Surface rendering of the bursts in C and D (insets) allowed the three-dimensional structure to be appreciated, demonstrating the complex architecture of the RNA signal. Although these two bursts contained multiple foci, other burst sites had more condensed USvRNA and appeared as single foci (data not shown), which is to be expected, given the stochastic nature of transcriptional bursting (11, 13, 57). This cell-to-cell heterogeneity suggests that RSV transcription sites are at different stages of the bursting cycle in each cell (and even within a single cell containing two integration sites), and the larger bursts are likely more active compared to the compact foci.

### Gag proximity to the transcriptional burst site did not enhance viral gene expression

The live cell imaging experiments shown in Figures 1-6 and Supplementary Movies used USvRNA and Gag protein altered by the insertion of exogenous tags to detect fluorescence signals. However, because such tags can affect RNA and protein trafficking, we performed quantitative analysis of images obtained using simultaneous immunofluorescence and smFISH in RSV-infected cells. Confocal z-stacks of cells were deconvolved and surfaces were generated using Imaris analysis software. The brightest USvRNA object(s) in each nucleus were considered to be transcriptional bursts, consistent with previous reports (11, 13).

We observed Gag at transcriptional burst sites in the nuclei of infected cells (Figure 9A) and found that most (77%) of the nuclear Gag condensates located nearest to an USvRNA burst were within a distance of 1 µm (Figure 9B). In some cases, multiple Gag condensates were located a similar distance from the same transcriptional burst site (Figure 3C; Supplementary Movie 3). Using the Imaris surface function, we compared the intensities of Gag condensates closest to USvRNA bursts to the intensities of Gag foci farther away from the bursts. The Gag condensates closest to the bursts (mean intensity 299.4 A.U. ± 34.96) were significantly brighter than those farthest from the bursts (198.9 A.U. ± 6.14 A.U.; p<0.0001) (Figure 9C). This finding is consistent with the observations from the live cell imaging experiments in which Gag condensates became brighter over time, but did not change in area (Figure 1). Interestingly, although the intensities of Gag foci closest to the burst were higher, there was no significant difference in the volumes of the foci, suggesting that the Gag condensates remained the same size regardless of their position, similar to the result found in our live cell experiments (Figures 1 and 9D).

In previously described cases of kissing between mRNA and transcriptional condensates, the close distance (<1 µm) between transcriptional condensates and the gene locus was associated with an increase in gene expression (11, 13, 59). Because we found that Gag condensates were close to USvRNA transcriptional burst sites (mean distance of 0.54 µm), we examined whether Gag altered viral transcriptional activity. Quantitative analysis revealed very low correlation between Gag proximity to the USvRNA burst and the volume (Figure 9E; r = -0.13) or intensity (Figure 9F; r =0.02) of the RNA focus, suggesting that Gag did not affect the level of USvRNA synthesis under these experimental conditions. These data indicate that the mechanism by which Gag is recruited to the USvRNA burst may not involve Gag interaction with an active transcriptional condensate. Gag may instead interact with a different host factor(s) for targeting to the active viral transcription site. These candidates may include members of the Mediator complex, transcription factors, splicing factors, and chromatin remodelers that we identified as potential Gag-interacting partners in our previous proteomic study (43). Further studies will be needed to assess whether Gag alters the activity of cellular genes, which was not tested in these experiments.

### Gag forms complexes with host transcriptional proteins in the nucleus

Mediator proteins and the transcription regulator CTCF are involved in the kissing interaction between *Sox2* mRNA and transcriptional condensates (11, 13). We identified several members of the Mediator complex, including Med26, in our RSV Gag proteome study, and follow up experiments revealed an interaction of Gag with Med26 in RSV infected cells (43). To examine whether Med26, CTCF, or the largest Mediator protein Med1, potentially play a role in targeting Gag to specific subnuclear sites, we used bimolecular fluorescence complementation (BiFC) (60-65). The rationale for this approach is based on the observation that Med26, Med1, and CTCF are ubiquitous in the nucleus and therefore, specific interactions are difficult to visualize. The use of BiFC requires close association of the two proteins being studied and allows for transient interactions to be observed because the reconstitution of the fluorophore is irreversible.

For the BiFC experiments, the N-terminal half of Venus fluorophore was fused to Med1 (pVN-Med1), Med26 (pVN-Med26), or CTCF (pCTCF-VN), and the Venus C-terminus was fused to Gag (pGag-VC) (Figure 10A). QT6 cells were transfected with 100 ng of each plasmid as indicated in Figure 10 and imaged at 10 and 16 hours post transfection. All images were captured at the same resolution, gain, and laser power to allow for intensity comparison. Histograms were normalized based on the VN-Med26 + Gag-VC signal because it was the brightest. In cases in which the signals for Med1 or CTCF were too low to see even when normalized, a second image at higher fluorescence intensity was shown with uniform adjustments for each image. Each plasmid was expressed alone did not fluoresce confirming the specificity of the interaction (Supplementary Figure 3) (24, 31).

**Figure 10:**
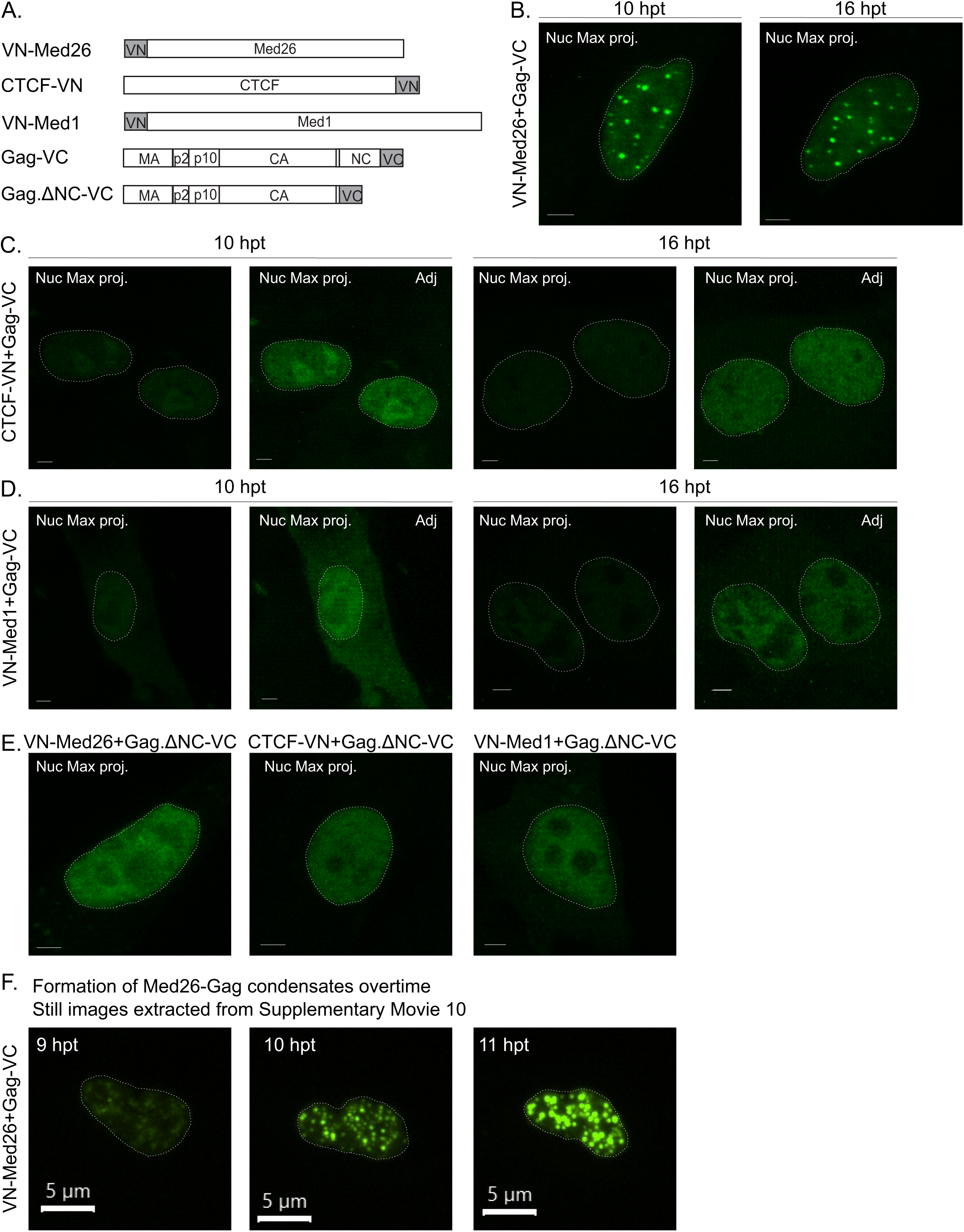
Gag forms complexes with Med26, CTCF, and Med1 in the nucleus as observed via Bimolecular fluorescence complementation (BiFC). **A)** Schematic of the constructs used in this complementation assay. The constructs are fused to either the N-terminus (VN) or C-terminus (VC) of Venus. Fluorescence is only produced when both halves come together to re-constitute the protein, indicating that the proteins the Venus halves are fused to are within close proximity. **B-E)** Cells were transfected with 100 ng of the indicated constructs and fixed at either 10 hours (3 replicates) or 16 hours (5 replicates) post-transfection (hpt). Z-stacks of nuclei were imaged via confocal microscopy and used to generate maximum projections. The nuclei are outlined in white. Scale bar = 2 µm. All cells were imaged exactly the same and histograms were adjusted in relation to the VN-Med26 + Gag-VC signal **(B)**. In the cases when the signal is very low, a second adjusted image is presented (Adj). **B)** When VN-Med26 and Gag-VC come into close proximity, in a majority of cells, diffuse and foci signal is present in the nucleus. **C)** CTCF-VN-Gag-VC complexes form diffuse signal in the nucleoplasm and nucleolus. **D)** Very few cells contained fluorescence signal when transfected with VN-Med1 and Gag-VC. Cells had diffuse cytoplasmic and nuclear signal. **E)** To determine if Gag nucleic acid binding ability is required for the interaction with Med26, CTCF, and Med1, a Gag.deltaNC-VC construct was used (2 replicates). Signal was diffuse in the nucleus in all instances. Cells with fluorescence signal when transfected with VN-Med1 and Gag.deltaNC-VC were more numerous than with wild-type Gag. **F)** To observe the formation of VN-Med26+Gag-VC BiFC signal over time. QT6 cells were transfected with 100 ng and imaged starting at 8 hpt. Stills correlating to the 9, 10, and 11 hpt from Supplementary movie 10 show the formation of foci and brighter signal over time. N=2 replicates. Scale bar = 5 µm.

In cells expressing VN-Med26 and Gag-VC, most cells contained focal and diffuse Venus fluorescence (Figure 10B) at each of the timepoints. To determine whether there was an increase in the number of foci and the signal intensity over time, live cell imaging was performed beginning at 8 hours post-transfection (hpt) and continuing for 3 hours with Z-stacks captured every 10 minutes. In images extracted from Supplementary Movie 10 (Figure 10F), we observed a marked increase in the number and size of Gag-Med26 foci and Venus fluorescence intensity as the imaging time progressed. These data suggest that the interaction between Gag and Med26 is a dynamic process and occurs after Gag has entered the nucleus with increased association over time. In addition, these Gag-Med26 foci appear to be BMCs, as there were numerous instances of fusion between neighboring condensates (Supplementary Figure 4).

In cells expressing CTCF-VN + Gag-VC, most cells contained diffuse nucleoplasmic signal with a subset of cells displaying nucleolar localization (Figure 10C), as reported previously for CTCF (66). For VN-Med1 + Gag-VC, the signal was diffuse throughout the nucleus, with a small amount of cytoplasmic signal observed, suggesting these proteins could interact initially in the cytoplasm prior to nuclear import (Figure 10D). As an indication of the strength of the interaction, many more cells exhibited fluorescence for Gag + Med26 and Gag + CTCF compared to Gag + Med1. The number of cells expressing fluorescence was quantitated, with 46 positive cells for Med 26 (average 2.3 cells/field for 20 fields), 49 positive cells for CTCF (average 2.5 cells/field for 20 fields), and very few positive cells for Med1, with only 6 total positive cells found in 5 experiments.

The NC (nucleocapsid) domain of Gag is responsible for Gag-Gag and Gag-nucleic acid interactions. Furthermore, deletion of NC or its replacement with a zip domain that allows for protein-protein but not protein-nucleic acid interactions prevents the formation of Gag nuclear foci, suggesting that formation of Gag nuclear foci depends on the RNA binding function of NC (31). Based upon this evidence, we performed BiFC with Gag.ΔNC-VC to determine whether the interactions between Gag and each cellular factor were NC-dependent (Figure 10E). Although fluorescence was observed between the NC-deleted Gag mutant and each cellular factor, the specificity of the interaction was lost. VN-Med26 + Gag.ΔNC-VC no longer formed foci. These data suggest that the formation of foci by the interaction of Gag with Med26 is NC dependent and could be explained by its RNA binding activity or the absence of the NC IDR, which may mediate formation of Gag-Med26 co-condensates (23, 67). With Gag.ΔNC, nucleolar localization of the Gag-CTCF-BFIC signal was lost, likely due to the deletion of the nucleolar localization signal in the NC domain (26). To our surprise, in the absence of NC, the number of cells with VN-Med1 + Gag.ΔNC-VC BiFC signal increased from 6 total cells in 5 experiments with WT Gag-VC (Figure 10D) to 11 total cells among 2 experiments (Figure 10E), providing further evidence that NC serves a regulatory role in the interactions of Gag with Med26, Med1, and CTCF.

Taken together, the data presented herein indicate that RSV Gag condensates enter the nucleus and interact with USvRNA burst sites co-transcriptionally through a dynamic kissing mechanism, in a similar fashion as transcriptional condensates with cellular genes (11, 13). In addition, the BiFC experimental results suggest that Gag traffics to these sites via interaction with Med26, CTCF, and/or another host factor. Furthermore, we presented evidence that the Gag-USvRNA complexes formed at sites of vRNA synthesis are subsequently exported from the nucleus, possibly for the purpose of encapsidation of gRNA into nascent virions.

## Discussion

Retroviruses cause severe disease including cancer and lethal immunodeficiencies, yet significant portions of the replication cycle remain poorly understood. Despite the absolute requirement for encapsidation of the viral genome for infectivity, it remains uncertain how retroviral Gag proteins find their RNA genomes for assembly into virions. Previously, it was shown that RSV Gag nuclear trafficking is required for efficient gRNA packaging, raising the possibility that recognition and capturing of gRNA occurs in the nucleus (24, 30). The Gag proteins of RSV and HIV-1 form BMCs, localize to viral transcription sites in the nucleus, and may interact with host transcription machinery, chromatin modulators, and splicing factors (24, 25, 35, 37, 40-43). Many key questions remain unanswered regarding how Gag condensates interact with cellular machinery to traffic to viral transcription sites, recognize and bind gRNA, and form vRNP complexes to nucleate assembly of virus particles (23-25, 35, 37, 43).

In the present study, we use live cell confocal microscopy and quantitative imaging analysis to gain insight into the mechanism by which the RSV Gag protein interacts with active viral RNA transcription sites. To our knowledge, the present study is the first to demonstrate a dynamic interaction of viral condensates with nucleic acids, as previous examples of kissing condensates involved cellular transcription clusters, enhancers, and mRNA synthesized at transcriptional bursts (11, 13). Our results suggest that retroviruses hijack mechanisms used by cellular condensates to traffic to viral transcription sites, interacting transiently with nascent RNA.

In live-cell experiments, we observed Gag condensates transiently co-localizing with nascent USvRNA at transcriptional bursts, presumably at sites of proviral DNA integration into the host chromosome. Kissing was defined as co-localization (distance of ≤0.25 µm) of the centers of Gag condensates with USvRNA foci (Supplementary Movies 1-4, Figures 1-3). We were able to capture a Gag condensate entering the nucleus close to the transcription site before co-localizing with the USvRNA burst (Supplementary Movie 5, Figure 4). Gag appeared to enter the nucleus within close proximity of the USvRNA raising the possibility that Gag could enter through specific nuclear pores (68, 69) or specialized regions of the nuclear envelope. In this study, we were unable to determine whether Gag displays directed movement or nuclear pore specificity. Thus, additional studies are needed to address these questions. Interestingly, we observed that Gag condensates became brighter over time even though the condensate area remained constant (Figure 1). Although this result suggests that more Gag molecules entered the condensate and became densely packed leading to an increase in brightness, it is unclear whether multiple Gag molecules bind the Ψ sequence or along downstream regions of the USvRNA. Future experiments will be needed to explore this question.

Experiments using dominant negative nucleoporin constructs containing the FG repeats from Nup98 and Nup214 allowed us to visualize Gag interactions with transcriptional bursts more frequently (Figure 6, Supplementary Movies 8 and 9) due to interference with the CRM1-mediated nucleocytoplasmic transport of Gag. Interestingly, in cells co-expressing NP98 or NP214, Gag appeared to be tethered with restricted movement, suggesting that activity of the dominant negative Nup mutants limits the transport of Gag from the transcription site to the nuclear envelope. Nup98 and Nup214 have functions in the nucleoplasm (70-74), in addition to being part of the nuclear pore complex, which may explain why interfering with their roles in transcription and chromatin remodeling could impair Gag mobility at transcription sites.

RSV integrates near the edge of the nucleus, as shown by our data indicating that USvRNA bursts localized within 1 µm of the nuclear rim (Figure 8). Finding that Gag also localizes near the nuclear periphery suggests that a mechanism similar to “gene gating” (75, 76) could facilitate Gag binding to active viral transcription sites near the nuclear envelope to shorten the journey into and out of the nucleus. Based on previous studies that HIV-1 Gag localizes within 1 µm of the edge of the nucleus and preferentially co-localizes with transcriptionally-active euchromatin marks (35), future studies will examine the interaction of RSV Gag with euchromatin histone marks near the nuclear periphery.

Using confocal microscopy, transcriptional bursts appeared as large bright foci, however, STED revealed that many bursts contained multiple small foci in clusters that could represent single RNAs (Supplementary Figure 2) at different stages of RNA synthesis, similar to super-resolution images of RNA polymerase II clusters (57). The dynamics of RNA bursting has been reported to be regulated by the proximity of transcription factors to the promoter, the number of transcription factor binding sites present, and their binding affinity (55). Transcription factors, including members of the Mediator complex, bind to clusters of enhancers and use the dynamic movement between the enhancer and promoter to interact with transcriptional condensates in a transient kissing interaction involving CTCF and cohesin to increase transcription (11). Based on those data, we speculate that chromatin looping is involved in the movement of Gag condensates toward the active viral transcription sites, and this possibility will be important to investigate in the future.

Our data revealed there was no correlation between Gag proximity to the viral transcriptional burst and viral transcriptional activity (Figure 9), suggesting that Gag does not modulate USvRNA synthesis. Because kissing interactions in the nucleus can occur within or between chromosomes to regulate gene expression (77, 78), it is possible that Gag takes advantage of these chromatin rearrangements to come into close proximity of the USvRNA burst. Furthermore, the RSV Gag interactome includes Mediator family members, RNAPII subunits, and splicing factors (37, 43) which may mediate the interaction of RSV Gag with vRNA-containing transcriptional condensates. Interestingly, the BiFC experiments reported herein revealed that Gag displayed a strong interaction with Med26 and forms nuclear foci resembling BMCs in an NC-dependent manner (Figure 10), suggesting that this association is mediated by NC through its nucleic acid binding ability or IDR-driven BMC formation. CTCF-Gag BiFC produced fluorescence with nucleolar localization that was dependent on the Gag NC domain. The Med1 interaction with Gag, however, increased when NC was deleted, suggesting that the NC domain plays a regulatory role in this interaction. Together, these data suggest that Med26 and CTCF interact with Gag in the nucleus to facilitate its localization to active sites of transcription.

We have wondered how Gag traffics specifically to sites of active viral transcription as opposed to sites of cellular gene expression. One possibility is that Gag moves toward the viral transcription site based on the RNA gradient established when viral gene expression is activated. The high affinity binding interaction between the Gag protein and psi-containing viral RNA could result in stable binding of Gag at the site of the emerging 5’ end of the USvRNA. Why is there a transient kissing interaction between Gag and USvRNA at the transcription site? Using computer modeling, it has been hypothesized that transcriptional condensates are attracted to the promotor due to the RNA gradient at the transcription site; low amounts of nascent RNA attracts the transcriptional condensate while high amounts of RNA repel the condensate, revealing how the kissing mechanism regulates transcriptional activity (79). We postulate that Gag condensates move toward low levels of transcribed viral RNA initially, Gag binds USvRNA at the burst, and as the RNA levels increase, vRNP co-condensates traffic away from the transcription site through the nuclear envelope, into the cytoplasm, and to the plasma membrane for virus assembly.

In our proposed model, Gag traffics into the nucleus, associates with euchromatin-bound proteins, and initially forms a co-condensate with Med26, CTCF, or other proteins that are distinct from a transcriptionally-active condensate (Figure 11, left side, non-colocalized). Other possible Gag-interacting host proteins that that could be involved in transport of Gag within the nucleus include splicing factors and nucleolar proteins, which also form BMCs (23, 26, 37, 43, 80). This interaction, along with the viral RNA gradient at the transcription site, may mediate Gag transport to the site of nascent USvRNA synthesis (Figure 11, right side, co-localized). We hypothesize that when the Gag condensate colocalizes with the USvRNA transcription site, it binds to nascent USvRNA and selects it for packaging. This hypothesis is compelling because co-transcriptional selection of gRNA by Gag would increase packaging efficiency, as the USvRNA captured by Gag would not become a spliced or full-length viral mRNA. Our previous work showed that Gag binding to the psi (Ψ) packaging sequence in the USvRNA causes a conformational change that exposes the nuclear export signal and enhances binding of the CRM1-RanGTP export complex, leading to Gag-USvRNA egress through the nuclear envelope (29) for the assembly of new virions at the plasma membrane.

**Figure 11:**
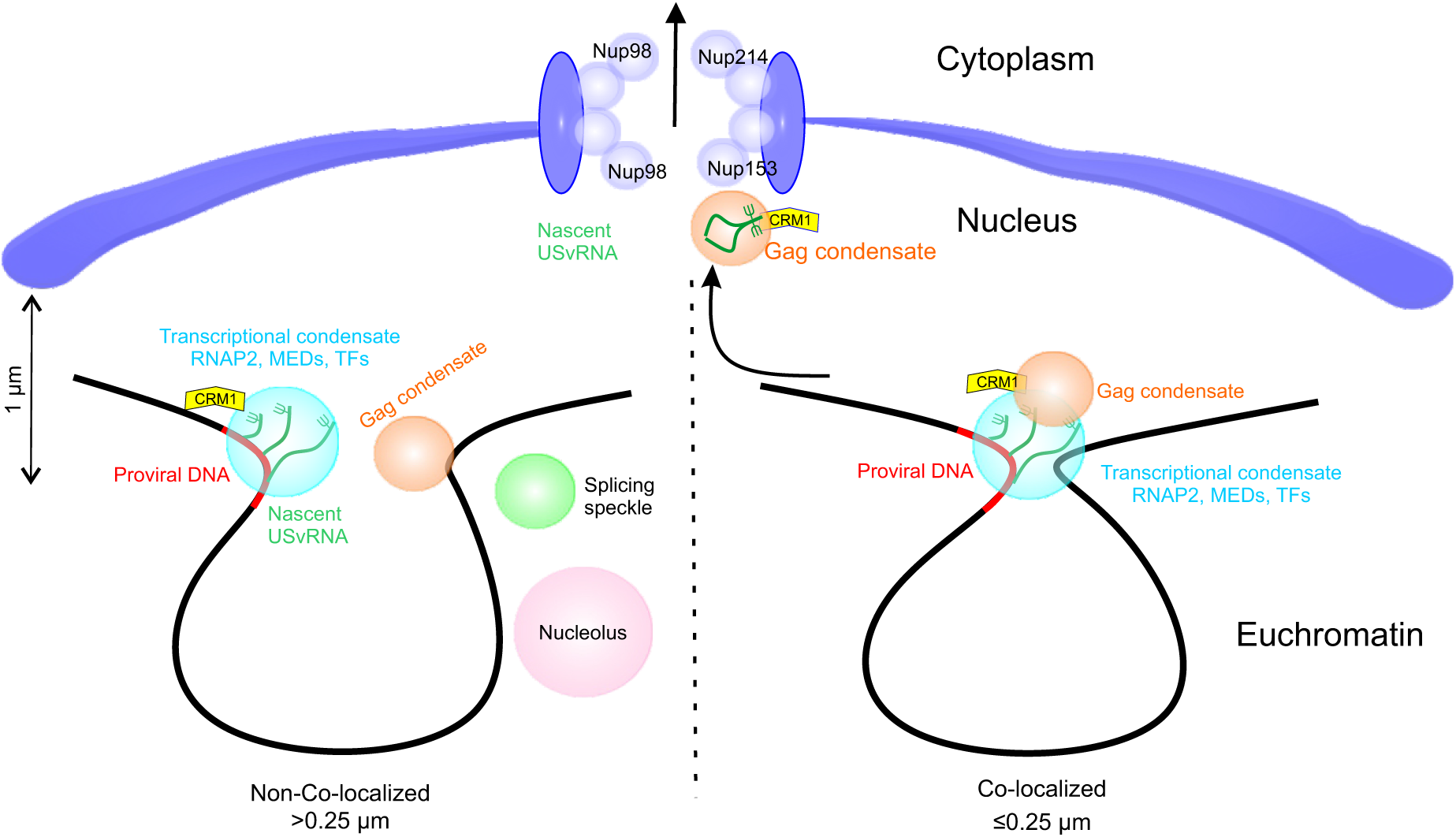
Model for Gag interaction with USvRNA at transcriptional bursts. In the nucleus, Gag binds to a cellular factor such as mediator proteins, transcription factors, splicing factors, chromatin, or nucleoli. When the condensate “kisses” the RSV integrated provirus gene locus where RNPII as part of the transcriptional condensate is transcribing USvRNA, Gag binds the USvRNA to form a viral ribonucleoprotein complex (vRNP). This complex is then exported from the nucleus through the nuclear pore via CRM1, traffics through the cytoplasm, and to the plasma membrane for virion assembly.

## Methods

### Plasmids and cell lines

Experiments were performed using chemically transformed QT6 quail fibroblast cells which were maintained and transfected via the calcium phosphate method as previously described (81-83).

Many of the constructs used to create the TRE RC.V8 constructs with internal tagged Gags and MS2 stem loops were based upon the cloning strategy used to clone pRC.V8 Gag-CFP 24xMS2 constructs, which was previously described (24). To create PB TRE RC.V8 MS2 stbl, first the region of RC.V8 encoding from the PmlI restriction site in *pol* to the end of the 3’LTR was amplified using primers 5’-TCTCCACGTGCGGAGTCATTCTGA-3’ and 5’-CGATGCGGCCGCCCCTCCGACGGTACTCAGCTTCTG-3’, and inserted into the PmlI and NotI sites of a piggybac TRE RC.V8 RU5 Gag.Pol mCherry construct with the first two ATG codons mutated to ATA to prevent translation (unpublished data) (PB TRE RC.V8 2ATG-ATA). To correct the ATA mutations to functional ATGs, PB TRE RC.V8 2ATG-ATA was digested with PmlI and NotI, and swapped with the corresponding sites in a PB TRE Gag-Pol plasmid containing functional ATGs to create PB TRE RC.V8. To insert 24 copies of MS2 stem loops, a restriction fragment from an RC.V8 derived construct that contained a stop codon after *nc* with 24 copies of MS2 stable stem loops between *nc* and *pr* were cloned into the FseI and PmlI sites of PB TRE RC.V8 to create the final PB TRE RC.V8 MS2 stbl construct. The location of the MS2 RNA stem loops between *nc* and *pr* allows for the specific labeling of unspliced viral RNA only by the MS2 coat protein. pCR4-24XMS2SL-stable was a gift from Robert Singer (Addgene plasmid # 31865; http://n2t.net/addgene:31865; RRID:Addgene_31865).

To create the PB TRE RC.V8 Gag-SNAPTag MS2 stbl construct, an RC.V8 derived construct that contained *gag-SNAPTag* and 24 copies of MS2 stem loops between *SNAPTag* and *pr* were cloned into the FseI and PmlI sites of PB TRE RC.V8 to create the final PB TRE RC.V8 Gag-SNAPTag MS2 stbl construct.

pSun1-Venus was created using Gibson assembly (84), with fragment 1 obtained by digesting pVenus-N2 with NheI and BamHI. The sequence encoding *sun1* (fragment 2) was amplified from pDEST-Sun1-mCherry (a gift from Jan Karlseder, Salk Institute for Biological Studies (85)) using primers 5’-ACCGTCAGATCCGCTAGCGCTATGGATTTTTCTCGGCTTCACATGTACAGT-3’ and 5’-CTCGCCCTTGCTCACGGATCCGGTGGCGACCGGTCCGATCA-3’, and was flanked by sequences that overlap the ends of fragment 1. pMS2-Halo-NLS was cloned by PCR amplifying the *halo* tag region from PB-H2B-Halo using primers 5’-ATCGACCGGTCGCCACCGGGATCCACGAAATCGGTACTGGCTTTCCATTCGACCCCCATT-3’ and 5’-CGATATCGATTTATACCTTTCTCTTCTTTTTTGGGGAAATCTCCAGAGTAGACAGCCAGC -3’, and inserted into the AgeI and ClaI restriction sites in pMS2-YFP-NLS (45, 46, 53). LZ10 PBREBAC-H2BHalo was a gift from James Zhe Liu (Addgene plasmid # 91564; http://n2t.net/addgene:91564; RRID:Addgene_91564) (86). The pGag-SNAPTag, NES1-YFP-MS2-NLS (a gift from Yaron Shav-Tal, Bar-Ilan University) (87), PB-t-rtTA, pSL-MS2-24x and pMS2-YFP-NLS, generous gifts from Dr. Robert Singer, Albert Einstein College of Medicine (45, 46), and pGag.L219A-CFP have been previously described (23-25, 88).

pVN-Med26 was cloned into pFLAG-Venus-Med26 (unpublished data) with the Med26 derived from FLAG-Med26, a gift from Joan Conaway & Ronald Conaway (Addgene plasmid# 15367; http://n2t.net/addgene:15367; RRID:Addgene_15367), via Gibson assembly (84, 89). pFLAG-Venus-Med26 was digested with SnaBI and NotI to remove FLAG and full-length Venus (Fragment 1). Part of the vector backbone was added back in by amplification from pFLAG-Venus-Med26 (Fragment 2) using primers: 5’-TTTCCTACTTGGCAGTACATCTAC-3’ and 5’-GCTAGCCAGCTTGGGTCTCCC-3’, and were flanked by regions overlapping fragments 1 and 3. VN was amplified from pSun1-Venus (Fragment 3) using primers: 5’-GGGAGACCCAAGCTGGCTAGCATGGGATCCGTGAGCAAGGGCGAG-3’ and 5’-GCCGGAGCCGCTGTGTGGATCGCGGCCGCCTCGATGTTGTGGCGGATCTTGAAGTT-3’, with overlaps complementary to the ends of fragments 1 and 2. Fragments 1 and 2 were the same from both pVN-Med26.

To create pCTCF-VN, VN was amplified from pSun1-Venus using primers: 5’-ATCGTACTTAAGGGGGGAGCAGGAGGCGGATCCGTGAGCAAGGGCGAG-3’ and 5’-TACGATGCTAGCGCGGCCGCTTACTCGATGTTGTGGCGGATCTT-3’. The VN PCR product was cloned into the AfIII and NheI sites in pKS070-pCAGGS-3XFLAG-(human)CTCF-eGFP to create the construct. pKS070 - pCAGGS-3XFLAG-(human)CTCF-eGFP was a gift from Elphege Nora (Addgene plasmid # 156448; http://n2t.net/addgene:156448; RRID:Addgene_156448)(90).

pVN-Med1 was cloned into a pFLAG-Halo-Med1 construct (unpublished) using Gibson assembly, with Med1 derived from pWZL hygro Flag HA TRAP220 wt, a gift from Kai Ge (Addgene plasmid # 17433; http://n2t.net/addgene:17433; RRID:Addgene_17433) (91). pFLAG-Halo-Med1 was digested with SnaBI and BstBI to create fragment 1. pVN-Med1 and fragment 2 was amplified from pFLAG-Halo-Med1 using primers: 5’-TTTCCTACTTGGCAGTACATCTAC-3’ and 5’-GCTAGCCAGCTTGGGTCTCCC -3’. VN (Fragment 3) was amplified from pSun1-Venus using primers: 5’-GGGAGACCCAAGCTGGCTAGCATGGGATCCGTGAGCAAGGGCGAG-3’ and 5’-TCCCCCTGAGCTTTCATATCGATTTCGAACCCCATATGTCCCAGGGAGGCATAATCAGGGACCTCGATGTTGTGGCGGATCTTGAAGTT-3’. Each fragment contained overlaps with the adjacent fragments. pGag-VC and pGag.ΔNC-VC were previously described (24, 31). pNP98 and pHA-NP214 Dominant Negative mutants were previously described (34).

To create the QT6 rtTA PB TRE RC.V8 MS2 stbl cell line, QT6 cells were seeded in a 35 mm dish at 0.3x10^6^ and transfected the next day with 3 µg of PB TRE RC.V8 MS2 stbl and 1.2 µg of transposase (System Biosciences) for a ratio of 0.2 µg of transposase per 500 ng of piggybac vector using the calcium phosphate method (24). Two days later the cells were transferred to a 100 mm dish. When the cells were ∼95% confluent, they underwent puromycin selection with 3 µg/mL of drug. Following testing and selection of the cell line, 1 µg of pPB-t-rtTA and 0.4 µg of transposase was transfected into the PB TRE RC.V8 MS2 stbl cell line. The QT6 rtTA PB TRE RC.V8 MS2 stbl cell line was subjected to selection with 2 µg/ml blasticidin. The QT6 rtTA PB TRE RC.V8 Gag-SNAPTag MS2 stbl cell line was created in the same fashion except the PB TRE RC.V8 Gag-SNAPTag MS2 stbl construct was transfected along with transposase using the same DNA amounts as before in a QT6 cell line that already expressed rtTA (QT6 rtTA).

### RC.V8 infection of QT6 cells

To create RC.V8-infected cells, uninfected QT6 cells were seeded into a 100 mm dish and the next day transfected with 10 µg of pRC.V8 via the calcium phosphate method. The next day, the media was changed. Virions were collected for ∼48 hours, centrifuged for 5 minutes at 2000 rpm at room temperature to remove dead cells, and added to naïve QT6 cells. Cells were infected at 37°C for 4 hours before changing the media. Cells were carried for prolonged periods.

### Simultaneous Immunofluorescence (IF) and smFISH

To visualize USvRNA and *cis*-expressed Gag in infected cells, cells were seeded at 0.5 x 10^6^ onto #1.5 coverslips. If cells were to be used for STED microscopy, they were transfected with 25 ng of pSun1-Venus via calcium phosphate to delineate the inner leaflet of the nuclear membrane for 16 hours. Cells were quick rinsed with RNase-free 1x PBS and fixed for 10 minutes in RNase-free 3.7% formaldehyde at room temperature, followed by 2x 5 minute washes with 1x PBS. The fixed cells were dehydrated in 70% ethanol at 4°C for a minimum of 24 hours. Cells were rehydrated in wash buffer (WB: 10% formamide, 2x SSPE, DEPC H_2_O) for 20 minutes at room temperature. Coverslips were incubated in a humid chamber for 16-20 hours at 37°C with 100 µl of hybridization buffer (10% dextran sulfate, 2x SSPE, 10% formamide) containing 1 µl of a 25 µM stock of 42 Stellaris RNA smFISH probes conjugated to Quasar 570 tiling the *gag* coding region (Biosearch) and mouse anti-RSV capsid primary antibody (made by Dr. Neil Christensen, Penn State College of Medicine) at 1:100. The next day, coverslips were incubated for 30 minutes at 37°C in WB containing donkey anti-mouse Alexa 647 (Thermo Fisher Scientific) at 1:1000. Coverslips were washed once more in WB for 30 minutes at 37°C either with (confocal) or without (STED) DAPI, and mounted in ProLong Diamond (Thermo Fisher Scientific).

### EU labeling of nascent RNAs

To visualize nascent RNAs, cells were pulse labeled with EU and labeled with Alexa 488 using the Molecular probes Click-IT RNA imaging kit. To visualize nascent RNAs in the QT6 rtTA TRE RC.V8 Gag-SNAPTag cell line, cells were seeded on coverslips as above and dox-induced for 24 hours. During the last 10 minutes, cells were pulse labeled with 1mM 5-ethynyl uridine (EU) at 37°C. Next, cells were rinsed 2x with 1x PBS, fixed for simultaneous IF to detect Gag and smFISH to detect USvRNA as above. Following 20 minutes of rehydration in WB, cells were rinsed 1x in 1x PBS and subjected to the Click-IT (click-chemistry, ThermoFisher Scientific) reaction to label the RNA with Alexa 488 for 30 minutes at room temperature. Coverslips were washed 1x in Click-IT rinse buffer and 1x in 1x PBS. The IF/FISH protocol was then completed as outlined above.

### Confocal Microscopy

For IF/FISH imaging, slides prepared as outlined above were imaged on a Leica AOBS SP8 FALCON confocal microscope equipped with hybrid detectors with time gating and a white light laser. Single fluorophore and secondary antibody controls were imaged to confirm that there was not any background or crosstalk. Slides were imaged with a 63x/NA 1.4 oil objective at a pixel format at 1024x1024, a scan speed of 400 Hz, and a 3x zoom. Z-stacks were captured at a step size of 0.3 µm with sequential scanning. Gag labeled via immunofluorescence was excited with a 647 nm laser line at 11% power and collected with a hybrid detector set to 652 nm-774 nm with a frame average of 2. USvRNA was excited with a 555 nm laser at 5% power and collected with a hybrid detector at 565 nm-630 nm with a frame average of 2.

The EU and Gag labeled QT6 rtTA TRE RC.V8 Gag-SNAPTag cell lines that were dox-induced for 48 hours were imaged similarly to infected cells except Gag-SNAPTag JF646 was excited with a 647 nm laser at 15% and collected from 652 nm-777 nm with a frame average of 4. USvRNA was excited with the 555 nm laser at 5% power and collected from 560 nm-630 nm with a frame average 4. EU Alexa 488-labeled RNA was excited with the 488 nm laser at 5% power and collected from 493 nm-540 nm with a frame average of 4. DAPI was excited with the 405 nm laser at 10% power and collected with a PMT with a frame average of 4.

For BiFC fixed cell imaging, QT6 cells were seeded were seeded onto coverslips at 0.4-0.5 x 10^6^ cells/coverslip. The next day, the cells were transfected with 100 ng of the desired BiFC plasmid and incubated for either 6, 10, or 16 hours. Cells were quick rinsed with RNase-free 1x PBS and fixed for 10 minutes in RNase-free 3.7% formaldehyde at room temperature, followed by 2x 5 minute washes with 1x PBS, and 1 minute DAPI stain. Coverslips were mounted on slides in Prolong Diamond and allowed to cure at room temperature for at least 24 hours. Slides were imaged on a Leica AOBS SP8 FALCON confocal microscope equipped with hybrid detectors with time gating and a white light laser. Single BiFC half controls were imaged to confirm that there was not any background. Slides were imaged with a 63x/NA 1.4 oil objective at a pixel format at 1024x1024, a scan speed of 400 Hz, and a 3x zoom. Z-stacks were captured at a step size of 0.3 µm with sequential scanning. BiFC signal was excited with a 514 nm laser line at 3% power and collected from 519 nm-587 nm. DAPI was excited with the 405 nm laser at 5% power and collected with a PMT. Images were processed in Imaris (Bitplane). Histograms were adjusted the same according to the VN-Med26 + Gag-VC signal for all images to show the differences in expression intensity. Some images are presented twice with further adjusted histograms for display

For live cell timelapse microscopy, QT6 rtTA TRE RC.V8 MS2 stbl cells were seeded onto glass bottom dishes (Mattek) at 0.5 x 10^6^ cells/dish. The next day, cells were transfected with 1 µg pNES1-YFP-MS2-NLS, and 500 ng of pGag-SNAPTag into transfection medium (5% fetal bovine serum (FBS) in DMEM) containing 2 µg/mL doxycycline to induce RC.V8 expression from the Tetracycline response element promotor (for 16+ hour inductions). One hour before imaging, cells were incubated with 50 nM of SNAPTag ligand JF549 [a kind gift from Luke Lavis, Janelia Research Campus (92)] for 1 hour at 37°C to label Gag-SNAPTag fusion proteins and either DRAQ5 or NucSpot650 (Biotium), where applicable. Cells were washed and imaged in imaging medium (clear DMEM with L-glutamine, 4.5 mg/liter D-glucose, 25 mM Hepes (Gibco) supplemented with 5% FBS, 9% tryptose phosphate broth and 1% chicken serum) at 16-22 hours post induction. For short term induction experiments (2 hours post induction), transfection media did not contain doxycycline and instead cells were doxycycline-induced for 1 hour before SNAPTag ligand labelling as above. Cells containing pNP98 or pHA-NP214 dominant negative FG repeats were transfected as above but also recieved1 µg of the respective NP construct. Cells were imaged between lines on a Leica AOBS SP8 FALCON confocal microscope in a live-cell incubated stage (Tokai Hit) at 37°C, 5% CO_2_ with a 63x/NA 1.2 water immersion objective at a rate of 1000 Hz at a frame every ∼1 s and a pixel size of 512 x 512. NES1-YFP-MS2-NLS was excited at 514 nm with 3% power and collected with a hybrid detector at 524 nm – 552 nm with time gating. Gag-SNAPTag JF549 was excited at 557 nm with 1% power and collected with a hybrid detector from 562 nm – 648 nm with time gating. NucSpot 650 or DRAQ5 live cell nuclear stain was excited with 653 nm at 3% laser power and collected with a PMT at 663 nm-779 nm.

For BiFC live cell imaging of pGag-VC and VN-Med26, QT6 cells were seeded onto glass bottom dishes (Mattek) at 0.5 x 10^6^ cells/dish. The next day, the cells were transfected with 100 ng of each plasmid and incubated for at least 6 hours. Cells were washed and imaged in imaging medium (clear DMEM with L-glutamine, 4.5 mg/liter D-glucose, 25 mM Hepes (Gibco) supplemented with 5% FBS, 9% tryptose phosphate broth and 1% chicken serum). Z-stacks (0.45 µm step size) were imaged every 10 minutes for 3 hours on a Nikon Ti2-E equipped with a Yokogawa CSU-X1 spinning disk, with a LUNF XL laser unit, Nikon Perfect Focus system, Z piezo stage, motorized XY stage, two sCMOS cameras (ORCA-Fusion BT, Hamamatsu Corp.). Cells were imaged with a 60x/NA 1.49 Apo TIRF oil objective with the correction collar optimized for 37°C and a live-cell incubated stage (Tokai Hit) at 37°C, 5% CO_2_. Images were collected in 12 bit with Standard mode and excited with the 488 nm laser with a 1 second exposure time. Images were cropped, histograms adjusted, and subjected to xy drift correction using Imaris (Bitplane).

For imaging Gag.L219A with nascent RNA in fixed cells, plasmids were transfected using the following ratio: 0.5 µg of MS2-YFP-NLS [a generous gift from Dr. Robert Singer, Albert Einstein College of Medicine] (45), 1.5 µg Gag.L219A-CFP and 3 µg of pSL-MS2-24x. Cells were fixed 12-24 hours post-transfection for 15 minutes with 3.7% paraformaldehyde in 2x PHEM buffer (3.6% PIPES, 1.3% HEPES, 0.76% EGTA, 0.198% MgSO_4_, pH to 7.0 with 10M KOH) (93), washed with 1x phosphate-buffered saline, DAPI stained, and mounted in anti-fade reagent (Invitrogen S2828). Slides were imaged using a Leica AOBS SP2 or SP8 confocal microscope, or a DeltaVision Elite Deconvolution microscope (GE).

### 5-Fluorouridine labeling of newly transcribed RNAs

QT6 cells were seeded onto coverslips as above and transfected with 1.5 µg of Gag.L219A-CFP using the calcium phosphate method. At 14-16 hours post transfection, cells were pulse labeled for 10 minutes with 2 mM 5-fluorouridine (5FU) in QT6 medium at 38.5°C and then fixed in 3.7% PFA in PHEM. Cells were permeabilized for 10-15 minutes at room temperature with 0.25% Triton X-100 in phosphate buffered saline (PBS) and blocked for 30 minutes in 10% bovine serum albumin (BSA) in PBS. RNAs incorporated with 5FU were detected with mouse anti-BrdU (Sigma B 2531) at 1:200 in 3% BSA/PBS for 1 hour at room temperature. Coverslips were incubated with either goat anti-Mouse Cy5 (Invitrogen A10524) or donkey anti-Mouse Alexa 647 (Life Technologies A31571) at 1:1000 for 1 hour at room temperature in 3% BSA/PBS, stained with DAPI, mounted, and imaged using a Leica AOBS SP8 confocal microscope or a DeltaVision Elite Deconvolution microscope (GE Heathcare Lifescience).

### Stimulated emission depletion (STED) super-resolution microscopy

For STED imaging of fixed cells, cells were prepared as above, without DAPI staining but with 25 ng of pSun1-Veus transfected to label the nuclear rim. Cells were imaged between lines on a Leica AOBS SP8 confocal microscope equipped with a STED module using a 100x/NA 1.4 oil immersion objective at 1000 Hz and a pixel format of 2048 x 2048. USvRNA was excited at 561 nm at 5% power and collected with a hybrid detector from 571 nm-620 nm, and depleted with the 775 nm laser at 50%. Sun1-Venus was excited with 514 nm at 6% laser power and collected with a hybrid detector from 524 nm-551 nm and depleted with the 592 nm laser at 30%. The Sun1 channel was also imaged with a frame accumulation of 2. All channels were imaged with Z STED at 50%.

For comparison between confocal and STED images of the USvRNA channel, the confocal channel was excited with the 561 nm laser at 10% power and collected with a hydrid detector at 571 nm-620 nm with a line accumulation of 2. The STED channel was excited and collected the same way except with depletion with the 660 nm laser at 50% and Z STED at 40%. Sun1-venus was imaged under confocal conditions. It was excited with a 514 nm laser at 10% power and collected with a hybrid detector at 524 nm – 541 nm with a line accumulation of 2.

### Quantitative image processing and data analysis

All confocal images and some STED images were deconvolved using Huygens Essential (SVI) using the classical maximum likelihood estimation (CMLE) deconvolution algorithm. Deconvolved z-stacks were further processed (Gaussian filters and histogram adjustments) and analyzed using Imaris image analysis 10.1.1 (Bitplane). The Imaris built-in machine learning algorithm was used to create surfaces of the DAPI (confocal), USvRNA, Gag, and EU channels. Any Gag, USvRNA, and EU surfaces outside of the nucleus were filtered out and removed from the analysis. Surface statistics were obtained including volume (µm^3^), sum signal intensity, distances between objects, and distance from the edge of the nucleus. The brightest RNA foci in each cell as determined by surface statistics were identified as transcriptional bursts (11, 13). Colocalization analysis of Gag.L219A-CFP with either pSL-MS2-24x or 5FU labeled nascent RNA was conducted as previously described (24).

Confocal and STED comparison images were deconvolved using the Huygens Essential low STED signal template. Surfaces of the Sun1 signal were created in Imaris using manual surface creation. Gag and RNA surfaces were created using machine-learning as above.

For live cell particle tracking, the Imaris spot function was used to identify Gag and USvRNA foci to determine the distance between Gag and the transcriptional burst over time. Also, the signal-based co-localization function in Imaris was used to generate a co-localization channel.

Graphs were generated and statistical analyses was performed in Prism (GraphPad) using an unpaired two-tailed *t* test. Outliers were identified and removed using a ROUT test, where appropriate. Pearson’s correlation (r) was used to determine the intensity correlations between Gag and USvRNA, and correlations between Gag distance to burst vs burst intensity/volume.

Four replicates (42 cells) were analyzed for IF/FISH confocal analysis. The BiFC experiments were conducted: 3x for 10 hpt, 5x 16 hpt, 2x for Gag. ΔNC-VC experiments. Live cell BiFC experiments were conducted 2x. For STED imaging of transcriptional bursts, three replicates were conducted, and 18 and 14 cells were imaged for STED alone and STED versus confocal analyses, respectively. 31 cells from 5 experiments were analyzed for Gag.L219A with pSL-MS2-24x and 18 cells from 3 experiments were analyzed for Gag.L219A with a 10 minute 5FU pulse. For live cell kissing experiments, for long dox induction 7 cells with kissing interactions were captured from 9 replicates. For short dox induction 4 cells from 6 replicates had kissing interactions. For live cell experiments with NP98, 10 cells with kissing events were captured from 2 replicates. In presence of NP214, 9 cells from 2 replicates had kissing events between Gag and USvRNA bursts.

## Acknowledgements

Luke Lavis (HHMI Janelia Research Campus) kindly provided the SNAPTag JF549 and JF646 ligands. Dr. Stephen Lockett (National Cancer Institute-Frederick National Lab for Cancer Research) wrote the MatLab colocalization scripts. We thank Gregory S. Lambert, Alexis Davison, Padmani Rai, Alecia M. Achimovich, and Jordan Chang, and Eunice C. Chen (Penn State College of Medicine) for critical discussions. We also thank Malgorzata Sudol for her work preparing the plasmids.

Microscopy images and were generated and processed in the Penn State College of Medicine Advanced Light Microscopy Core (RRID: SCR_022526). The Advanced Light Microscopy Core services and instruments used in this project were funded, in part, by the Pennsylvania State University College of Medicine via the Office of the Vice Dean of Research and Graduate Students and the Pennsylvania Department of Health using Tobacco Settlement Funds (CURE). The content is solely the responsibility of the authors and does not necessarily represent the views of the University or College of Medicine. The Pennsylvania Department of Health specifically disclaims responsibility for any analyses, interpretations or conclusions.

## Author Contributions

Conceptualization: R.J.K.M and L.J.P. Experimentation and Data analysis: R.J.K.M. Writing and Editing: R.J.K.M and L.J.P.

## Correspondence and request for materials

Correspondence can be addressed to either Rebecca J. Kaddis Maldonado or Leslie J. Parent. Request for materials can be address to Leslie J. Parent.

---

**Supplementary Figure 1:**
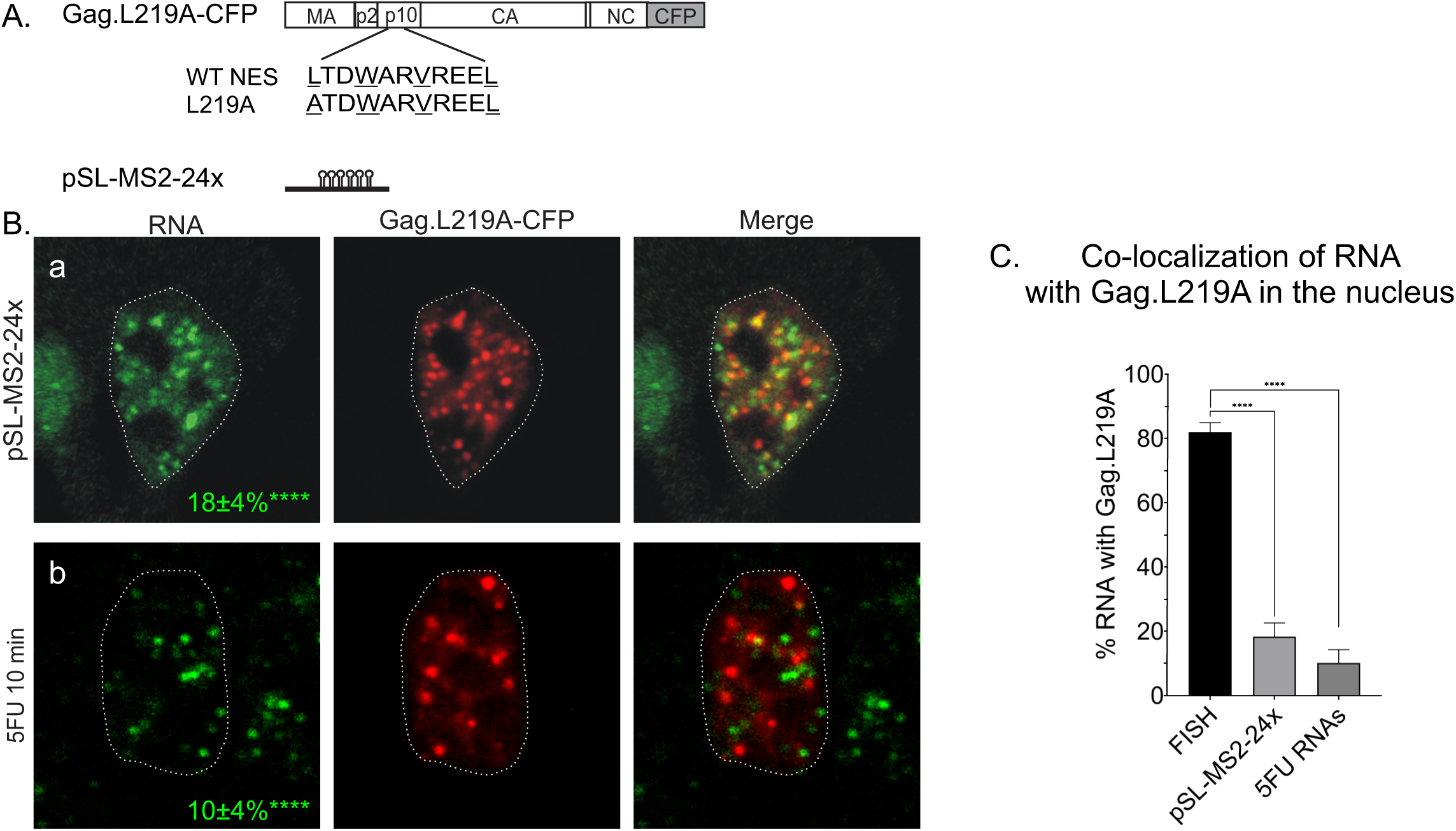
Co-localization between Gag.L219A and nascent non-viral RNAs. **A)** Schematic of the Gag.L219A-CFP NES mutant and pSL-MS2-24x. **B)** QT6 cells expressing a Gag nuclear export mutant (Gag.L219A) were co-transfected with either pSL-MS2-24x, a non-viral construct containing 24 copies of MS2 stem loops or pulse labeled for 10 minutes with 5-fluorouridine (5FU) to label nascent RNAs. (a) pSL-MS2-24x RNA foci (green) co-localized with Gag.L219A (red) at 18±4% (p<0.0001) in the nucleus (white outline). (b) 5FU labeled RNA foci (green) co-localized with Gag.L219A foci at 10±4% p<0.0001). **C)** Bar graph of percent co-localization of RNA with Gag.L219A (24). 31 cells from 5 replicates were collected for Gag.L219A+ pSL-MS2-24x and 18 cells from 3 replicates were analyzed for Gag.L219A + 5FU-labeled RNA.

**Supplementary Figure 2:**
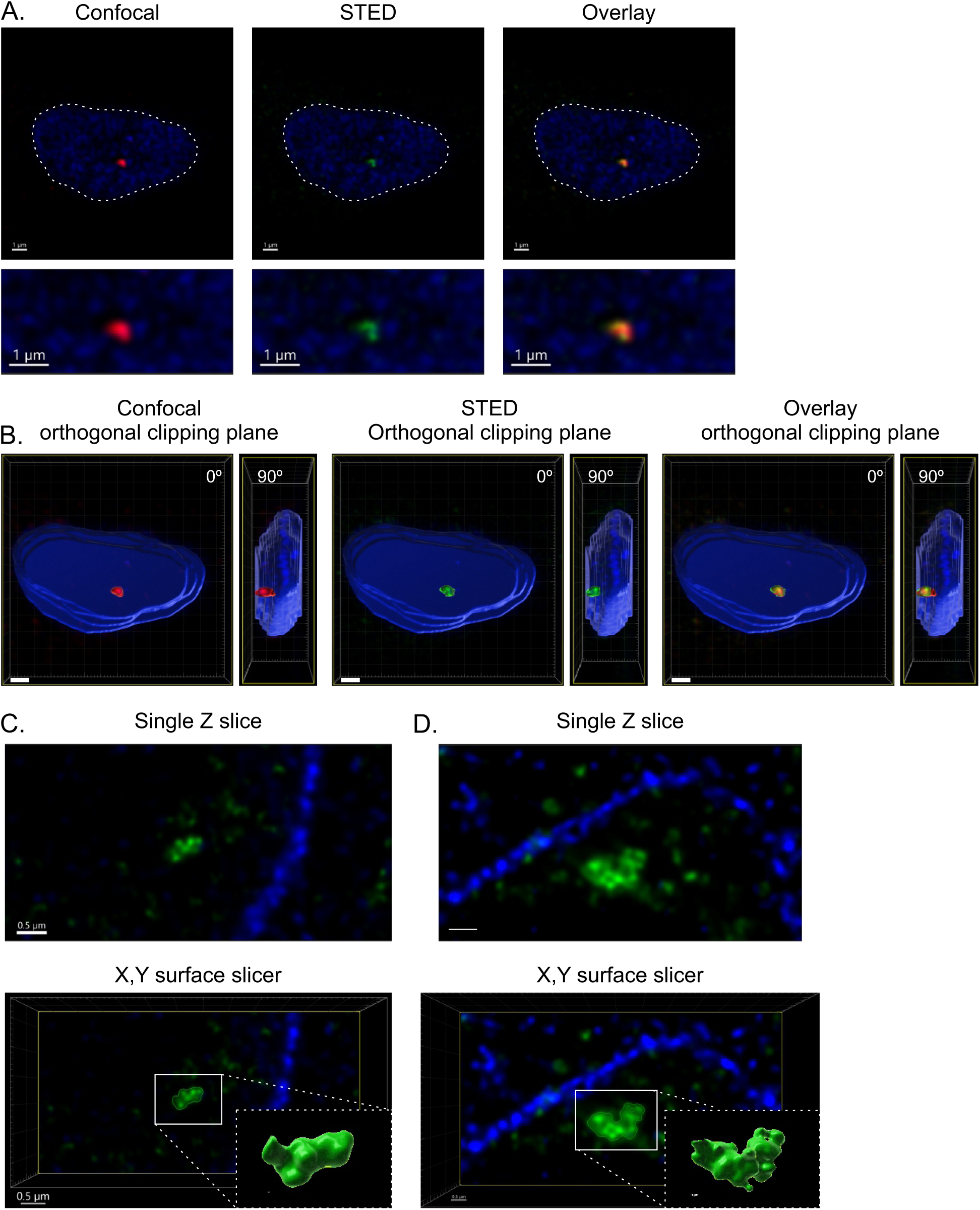
Single plasmid controls for BiFC. QT6 cells were transfected with VN-Med26, CTCF-VN, or VN-Med1 alone to confirm that fluorescence did not occur in the absence of Gag-VC. Cells were imaged a low power and adjusted according to VN-Med26+Gag-VC as for the images in figure 10. Scale bar = 10 µm. Six replicates were collected for each condition.

**Supplementary Figure 3:**
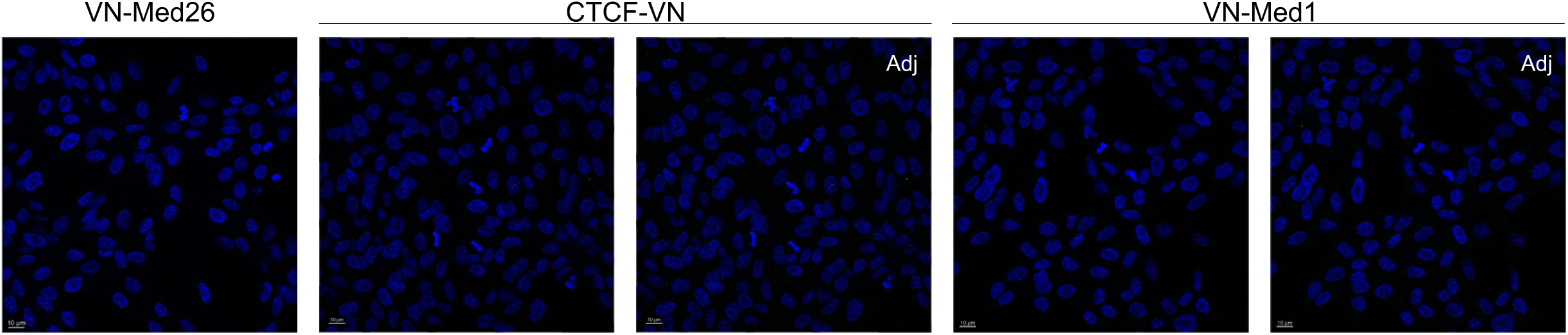
STED microscopy of USvRNA bursts reveals complex structures. **A)** Single z-slices of chronically infected cells comparing bursts of transcription imaged via confocal microscopy (red) to those imaged via STED (green). The nucleus is marked with Sun1-venus (blue, white outline). The image below is a zoom in of the burst of interest. The confocal burst appears as a single focus while the STED burst contains multiple smaller foci. Scale bar= 1 µm. **B)** Surface renderings were generated of the cell above and subjected to orthogonal clipping planes at either 0° or 90°. The STED bursts have a more lobed appearance. Scale bar= 1 µm. **C and D)** Two more examples of highly structured USvRNA bursts imaged via STED. The bursts are presented as a single Z-slice (Scale bar= 0.5 µm) or with an X,Y surface slicer (Scale bar=0.3-0.5 µm). In the bottom right corner of the bottom panels, a zoom in of a volume rendering of the bursts are presented (Scale bar= 0.07-0.1 µm). Both bursts appear lobed and highly structured. 3 Replicates.

**Supplementary Figure 4:**
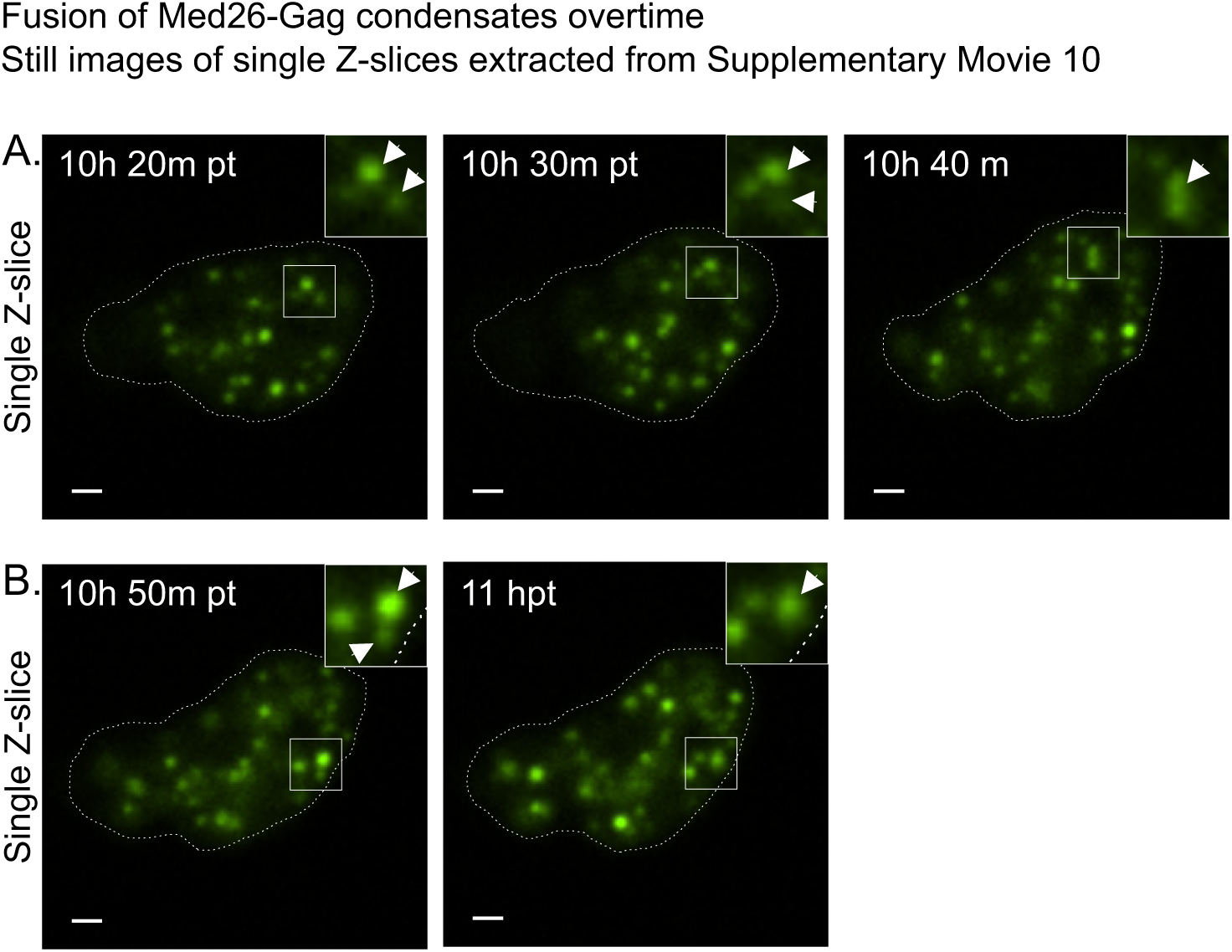
Fusion of VN-Med26:Gag-VC BMCs over time. Single Z slices were obtained from stills extracted from Supplementary Movie 10. Fusion of foci is a characteristic of BMCs. **A)** An example of two VN-Med26:Gag-VC foci (white arrows, inset) fusing over a period of 20 minutes. **B)** Another example of two VN-Med26:Gag-VC foci (white arrows, inset) fusing over a period of 10 minutes. Scale bar = 1 µm. 2 replicates

**Supplementary Movie 1:** The QT6 rtTA TRE RC.V8 MS2 stbl cell line that was transfected with Gag-SNAPTag JF549 (red) and NES1-YFP-MS2-NLS (USvRNA-green) and dox induced for approximately 22 hours. Cells were imaged every second. Particle tracking was conducted using the Imaris spot function. The USvRNA burst (green) and Gag focus (red) transiently co-localized in the nucleus. The nucleus is marked due to the NES1-YFP-MS2-NLS being able to clear the nucleus.

**Supplementary Movie 2:** This channel shows the overlay of the co-localization channel generated from the USvRNA and Gag signals from the movie presented in 1.

**Supplementary Movie 3:** The QT6 rtTA TRE RC.V8 MS2 stbl cell line was transfected with Gag-SNAPTag JF549 (red) and NES1-YFP-MS2-NLS (USvRNA-green) and dox induced for ∼16 hours. Cells were imaged every second. Particle tracking was conducted using the Imaris spot function. A burst of USvRNA (green) in the nucleus was met by a red focus of Gag to undergo a transient interaction. The nucleus was marked based on the NES1-YFP-MS2-NLS signal.

**Supplementary Movie 4:** The QT6 rtTA TRE RC.V8 MS2 stbl cell line that was transfected with Gag-SNAPTag JF549 (red) and NES1-YFP-MS2-NLS (USvRNA-green) and dox induced for ∼16 hours. Cells were imaged every second. Particle tracking was conducted using the Imaris spot function. Two Gag foci (red) were tracked to the same burst of USvRNA (green). Gag condensate 1: Yellow track. Gag condensate 2: White track. The nucleus is marked based on the NES1-YFP-MS2-NLS signal. This is the same cell imaged in Supplemental Movie 3 but at an earlier time point.

**Supplementary Movie 5:** The QT6 rtTA TRE RC.V8 MS2 stbl cell line that was transfected with Gag-SNAPTag JF549 (red) and NES1-YFP-MS2-NLS (USvRNA-green) and dox induced for 2 hours. Cells were imaged every second. Particle tracking was conducted using the Imaris spot function. A focus of Gag (red) was tracked from the cytoplasm into the nucleus and kissed the USvRNA burst (green). The nucleus is marked due to the NES1-YFP-MS2-NLS being able to clear the nucleus.

**Supplementary Movie 6:** The QT6 rtTA TRE RC.V8 MS2 stbl cell line was transfected with Gag-SNAPTag JF549 (red) and NES1-YFP-MS2-NLS (USvRNA-green) and dox induced for ∼22 hours. Cells were imaged every second. Particle tracking was conducted using the Imaris spot function. A focus USvRNA (green), not correlating to a burst, in the nucleus formed a vRNP with Gag that trafficked from the nucleus into the cytoplasm. The nucleus was labeled with NucSpot 650.

**Supplementary Movie 7:** This channel shows the tracking of the co-localization channel generated from the USvRNA and Gag signals from the movie presented in 6.

**Supplementary Movie 8:** The QT6 rtTA TRE RC.V8 MS2 stbl cell line that was transfected with Gag-SNAPTag JF549 (red), NES1-YFP-MS2-NLS (USvRNA-green), and NP98 (to trap Gag in the nucleus), and dox induced for 2 hours. Cells were imaged every second. Particle tracking was conducted using the Imaris spot function. A Gag condensate (red) kissed the USvRNA burst (green). The nucleus is marked via DRAQ5. A co-localization channel (white) was overlaid with the Gag and USvRNA signals to indicate when the condensates “kiss” over time.

**Supplementary Movie 9:** The QT6 rtTA TRE RC.V8 MS2 stbl cell line that was transfected with Gag-SNAPTag JF549 (red), NES1-YFP-MS2-NLS (USvRNA-green), and NP214 (to trap Gag in the nucleus), and dox induced for 2 hours. Cells were imaged every second. Particle tracking was conducted using the Imaris spot function. Three Gag condensates (red) kissed the USvRNA burst (green). The nucleus is marked via DRAQ5. A co-localization channel (white) was overlaid with the Gag and USvRNA signals to indicate when the condensates “kiss” over time.

**Supplementary Movie 10:** QT6 cells were transfected with VN-Med26 and Gag-VC. Fluorescence is only visible if Med26 and Gag come into close enough proximity to reconstitute the VN and VC halves into a full Venus protein. Cells were imaged starting at 8 hours post transfection and imaged every 10 minutes for 3 hours. Med26-Gag complexes form over time indicated by the increasing fluorescence and foci formation.

