## Supplementary Table 1 for "Dynamic interactions of retroviral Gag condensates with nascent viral RNA at transcriptional burst sites: implications for genomic RNA packaging"

| Supplementary Table 1: Burst distance to DAPI Edge | | | | | | | | | | |
|---|---|---|---|---|---|---|---|---|---|---|
| **Bin Center Distance From DAPI edge (µm)** | **0.00** | **0.1** | **0.2** | **0.3** | **0.4** | **0.5** | **0.6** | **0.7** | **0.8** | **0.9** |
| **Number of Bursts** | 27 | 8 | 13 | 5 | 4 | 12 | 14 | 4 | 6 | 2 |
