## Supplementary Table 2 for "Dynamic interactions of retroviral Gag condensates with nascent viral RNA at transcriptional burst sites: implications for genomic RNA packaging"

| Supplementary Table 2: Gag distance to DAPI Edge | | | | | | | | | | | | | | | |
|---|---|---|---|---|---|---|---|---|---|---|---|---|---|---|---|
| **Bin Number Distance to DAPI Edge (µm)** | **0.00** | **0.10** | **0.20** | **0.30** | **0.40** | **0.50** | **0.60** | **0.70** | **0.80** | **0.90** | **1.00** | **1.10** | **1.20** | **1.30** | **1.40** |
| Number of Gag foci | 822 | 132 | 100 | 102 | 46 | 30 | 28 | 12 | 15 | 17 | 9 | 6 | 5 | 0 | 4 |

| | | | |
|---|---|---|---|
| **1.50** | **1.60** | **1.70** | **1.80** |
| 1 | 1 | 0 | 1 |
